# A deep-sea sulfate reducing bacterium directs the formation of zero-valent sulfur via sulfide oxidation

**DOI:** 10.1101/2021.03.23.436689

**Authors:** Rui Liu, Yeqi Shan, Shichuan Xi, Xin Zhang, Chaomin Sun

## Abstract

Zero-valent sulfur (ZVS) is a critical intermediate in the biogeochemical sulfur cycle. Up to date, sulfur oxidizing bacteria have been demonstrated to dominate the formation of ZVS. In contrast, formation of ZVS mediated by sulfate reducing bacteria (SRB) has been rarely reported. Here, we report for the first time that a typical sulfate reducing bacterium *Desulfovibrio marinus* CS1 directs the formation of ZVS via sulfide oxidation. In combination with proteomic analysis and protein activity assays, thiosulfate reductase (PhsA) and sulfide: quinone oxidoreductase (SQR) were demonstrated to play key roles in driving ZVS formation. In this process, PhsA catalyzed thiosulfate to form sulfide, which was then oxidized by SQR to form ZVS. Consistently, the expressions of PhsA and SQR were significantly up-regulated in strain CS1 when cultured in the deep-sea cold seep, strongly indicating strain CS1 might form ZVS in its real inhabiting niches. Notably, homologs of *phsA* and *sqr* widely distributed in the metagenomes of deep-sea SRB. Given the high abundance of SRB in cold seeps, it is reasonable to propose that SRB might greatly contribute to the formation of ZVS in the deep-sea environments. Our findings add a new aspect to the current understanding of the source of ZVS.

## Introduction

Zero-valent sulfur (ZVS) is a central intermediate in the biogeochemical sulfur cycle^1–3^, and forms conspicuous accumulations at sediment surface under the sea floor including the cold seep and the hydrothermal systems^4–6^. In marine environments, ZVS commonly occurs in some forms such as polysulfides (S_n_), polymeric sulfur (S_n_) or cyclooctasulfur (S_8_)^7, 8^. The production of ZVS has been regarded as a bio-signature of sulfur-oxidizing microorganisms^9, 10^. The process of ZVS production begins with the formation of polysulfide through the oxidation of thiosulfate or sulfide^3, 11–13^. For the formation process of ZVS mediated by thiosulfate oxidation, there are at least four pathways identified in the sulfur oxidizing bacteria (SOB)^11, 13^, including Sox pathway^14^, tetrathionate (S_4_I) intermediate pathway^15^, Sox-S_4_I interaction system^16^ and a novel pathway mediated by thiosulfate dehydrogenase (TsdA) and thiosulfohydrolase (SoxB)^17^. For the formation process of ZVS mediated by sulfide oxidation, sulfide: quinone oxidoreductase (SQR) has been proposed to be the key enzyme to catalyze the formation of ZVS in various sulfur-oxidizing Alphaproteobacteria and Gammaproteobacteria^11, 12, 18^. SQR is a membrane associated protein that oxidizes sulfide to ZVS and transfers electrons to the membrane quinone pool with flavin adenine dinucleotide^19^. As a key sulfide detoxifying enzyme, SQR is present in many bacteria, archaea and the mitochondria of eukaryotic cells, classified into six types (Type I to VI)^19–21^. Since no more pure cultures containing the *sqr* gene are available^22, 23^, the function of SQR in *D*-proteobacteria (e.g. most of typical sulfate reducing bacteria) is obscure.

Up to date, most of progresses about the formation of ZVS are related to sulfur oxidizing bacteria but rarely associated with sulfate reducing bacteria (SRB). Notably, a novel pathway of ZVS generation mediated by the dissimilatory sulfate reduction has been recently observed in a syntrophic consortium of anaerobic methanotrophic archaea (ANME) and SRB^24–26^. ANME were firstly proposed to drive the formation of ZVS via coupling the anaerobic methane oxidation (AOM) with the sulfate reduction^25^, which provided experimental evidence for the first time to confirm the ZVS generation from the dissimilatory sulfate reduction. However, this proposal was challenged by other researchers, who insisted that the passage of sulfur species by ANME as metabolic intermediates for their SRB partners was unlikely^27^. In addition, based on a methanogenic bioreactor system and metagenomics approaches, some researchers proposed a novel ZVS formation pathway mediated by dissimilatory sulfate reduction^24, 28^, in which SRB might utilize sulfate-to-ZVS as an alternative pathway to sulfate-to-sulfide to alleviate the inhibitive effects of sulfide. This proposal was also needed to be further verified given that the canonical pathway of the dissimilatory sulfate reduction mediated by SRB reduces sulfate to sulfide without the production of ZVS^29, 30^. Till to date, the pure culture of SRB has not been isolated from both AOM enrichment cultures and the methanogenic bioreactor as mentioned above^25, 28^, which hindering the researchers to test whether SRB could directly drive ZVS formation via dissimilatory sulfate reduction or some other unknown pathways.

The typical dissimilatory sulfate reduction system contains a combination of sulfate adenylyltransferase (Sat) and adenylyl-sulfate reductase (AprAB) initiating the reduction of sulfate to sulfite, and then sulfite reductases catalyzing the reduction of sulfite to sulfide^31^. On the other hand, sulfide could also be produced by thiosulfate disproportionation process in SRB^32^. Potentially, SRB might form ZVS or even elemental sulfur from sulfide driven by SQR or other proteins with similar functions. However, to our best knowledge, there is no evidence that pure isolate of SRB could form ZVS via oxidizing sulfide that generated by dissimilatory sulfate reduction or thiosulfate disproportionation.

In the present study, a strictly anaerobic strain of *Desulfovibrio marinus* CS1 was isolated from the surface sediments of a cold seep in the South China Sea, and it was surprisingly found to form ZVS in the presence of thiosulfate. In combination with genomic, proteomic and biochemical approaches, PhsA and SQR were demonstrated to be responsible for the formation of ZVS in strain CS1, which is a novel pathway driving ZVS formation present in SRB. Based on the metagenomics analysis, the broad distribution of this novel pathway and its potential contribution to the deep-sea sulfur cycle were also investigated and discussed.

## Results

### Cultivation and identification of a typical sulfate reducing bacterium *Desulfovibrio marinus* CS1 from the deep-sea cold seep

Despite a high proportion of SRB has been reported in deep-sea cold seeps^33–36^, the lack of cultured representatives from deep sea has hampered a more detailed exploration of this important group. With this, we anaerobically enriched the surface sediment samples collected from the deep-sea cold seep with a modified sulfate reducing medium (SRM) at 28 °C for one month, enriched samples were then plated on the solid SRM in Hungate tubes. Given the presence of Fe^2+^ in the medium, typical SRB would form black FeS precipitation. As expected, the enrichment formed a large amount of black color colonies in the solid SRM, indicating the dominant presence of SRB in the enrichment. Single colonies with black color were subsequently purified several times using the dilution-to-extinction technique at 28 °C under a strict anaerobic condition. Surprisingly, based on the 16S rRNA gene sequencing, these colonies belong to a same SRB strain designated CS1. The 16S rRNA gene sequence of strain CS1 shared a high similarity of 99.54% with *Desulfovibrio marinus* E-2^T^ (accession no. NR_043757.1). Additionally, strain CS1 also clustered with *D. marinus* E-2^T^ according to the phylogenetic analysis (Supplementary Fig. 1a). The ANI and AAI analyses of genomes between strain CS1 and another strain *D. marinus* P48SEP (accession no. ASM762508) were respective 98.95% and 99.87% (Supplementary Figs. 1b and 1c), which were higher than the accepted threshold (both ANI and AAI value above 95%) for same species^37, 38^. Thus, strain CS1 was identified as a member of *D. marinus* and designated *D. marinus* CS1 in this study. Accordingly, *D. marinus* CS1 showed black color in the agar plate containing Fe^2+^ (Supplementary Fig. 2a). Under transmission and scanning electron microscopy observation, the cells of strain CS1 were short rod-like, approximately 2 μm × 0.5 μm in size, and had a single flagellum (Supplementary Figs. 2b and 2c).

### Diverse sulfur metabolic pathways existing in *D. marinus* CS1

To obtain a deeper insight into the characterization of *D. marinus* CS1, the genome of *D. marinus* CS1 was completely sequenced (Supplementary Table 1). When analyzing the genome sequence of strain CS1, we found a complete dissimilatory sulfate reduction pathway and a partial assimilatory sulfate reduction pathway present in *D*. *marinus* CS1 (Supplementary Fig. 2d, Supplementary Table 2). For dissimilatory sulfate reduction pathway, two conserved gene clusters responsible for transforming sulfate to sulfite and sulfite to sulfide (Supplementary Fig. 2e), were identified in *D*. *marinus* CS1. Notably, a homologous gene encoding SQR that usually involved in sulfide oxidation was surprised to be identified in the genome of *D*. *marinus* CS1 (Supplementary Fig. 2d and Supplementary Table 2). Given the presence of SQR-like protein and its usual sulfur oxidation activity, we speculated that some unexplored sulfur oxidation pathways might exist in strain CS1.

### Responses of *D. marinus* CS1 to different sulfur sources

Considering the presence of diverse sulfur metabolic pathways in *D. marinus* CS1, we sought to explore the responses of strain CS1 to different sulfur containing compounds including sulfate, sulfite, thiosulfate and sulfide. First, we monitored the growth dynamics of strain CS1 that cultured in SRM medium supplemented with above sulfur sources for up to two months. Surprisingly, the supplement of different sulfur sources showed a similar or even better growth status compared to the control group when cultured strain CS1 for up to two months, though some of them inhibited bacterial growth at the beginning of the incubation period (Fig. 1a). Especially, the supplement of 40 mM Na_2_S_2_O_3_ or 10 mM Na_2_SO_3_ in the medium could significantly promote the growth of strain CS1 when the cells entered the stationary phase (Fig. 1a). Therefore, strain CS1 showed different responses to different sulfur sources. On the other hand, cells of strain CS1 showed an extended morphology under the treatment of Na_2_S_2_O_3_ (Fig. 1d), Na_2_SO_3_ (Fig. 1e) and Na_2_S (Fig. 1f) compared to that in the control group (Fig. 1a) or supplement of Na_2_SO_4_ (Fig. 1b).

**Fig. 1.**
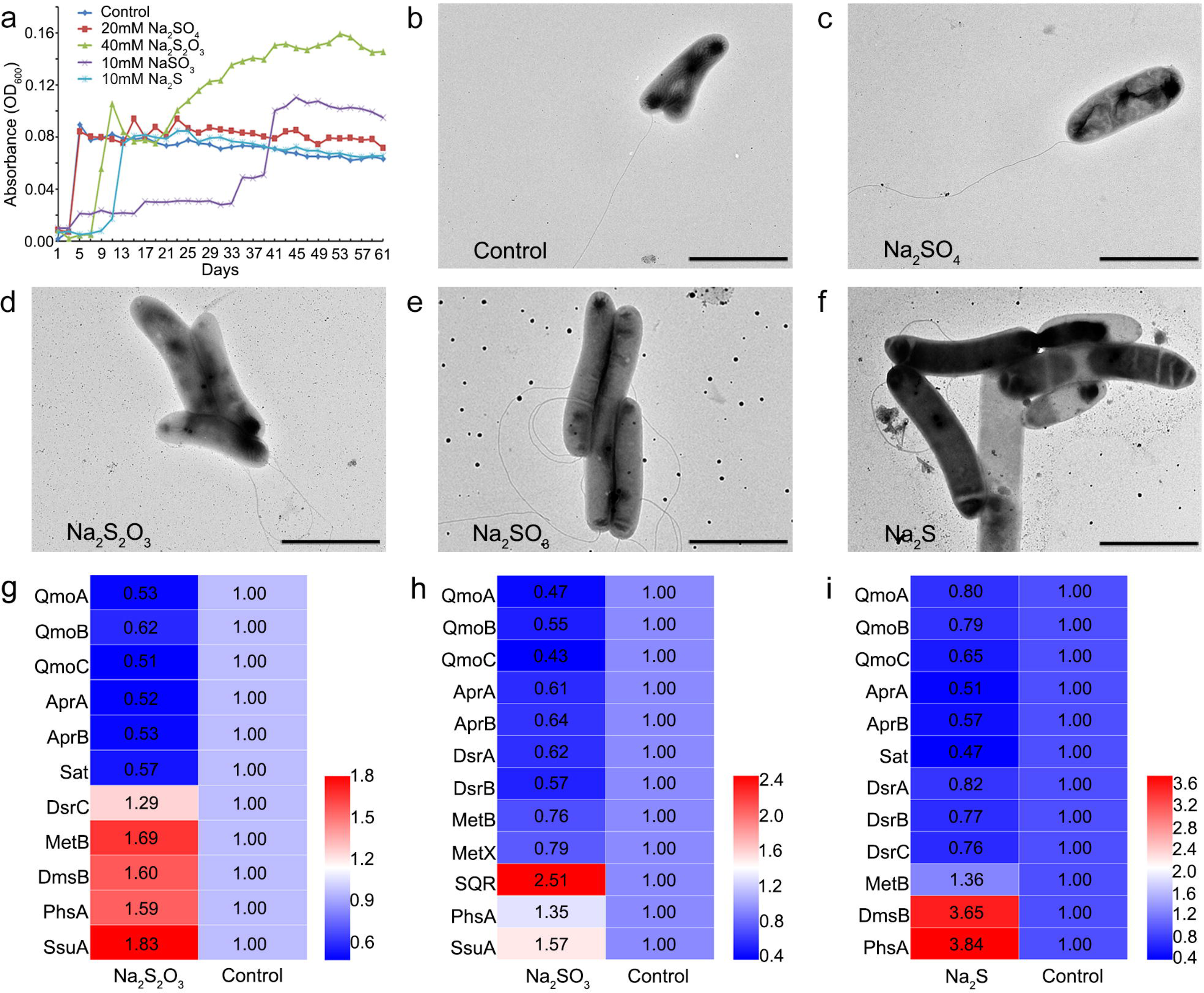
Responses of *D*. *marinus* CS1 to different sulfur sources. (a) Growth assays of *D*. *marinus* CS1 that cultured in the SRM medium supplemented with different sulfur sources, including Na_2_SO_4_ (20 mM), Na_2_S_2_O_3_ (40 mM), Na_2_SO_3_ (10 mM) and Na_2_S (10 mM). “Control” indicates *D*. *marinus* CS1 was cultured in the SRM medium. (b-f) TEM observation of morphology changes of *D*. *marinus* CS1 that cultured in the medium supplemented with different sulfur sources as shown in panel A. Scale bars, 2 μm. (g-i) Proteomics based heat map showing all significantly down- and up-regulated proteins associated with sulfur metabolism in *D*. *marinus* CS1 when cultured in the medium supplemented with 40 mM Na_2_S_2_O_3_ (g), 10 mM Na_2_SO_3_ (h) and 10 mM Na_2_S (i), respectively. The numbers shown in the heat map represent the fold change of proteins compared to the control group. Abbreviations: QmoA, CoB-CoM heterodisulfide reductase iron-sulfur subunit A family protein; QmoB, hydrogenase iron-sulfur subunit; QmoC, quinone-interacting membrane-bound oxidoreductase complex subunit QmoC; AprA, adenylyl-sulfate reductase subunit alpha; AprB, adenylyl-sulfate reductase subunit beta; Sat, sulfate adenylyltransferase; DsrA, dissimilatory-type sulfite reductase subunit alpha; DsrB, dissimilatory-type sulfite reductase subunit beta; DsrC, TusE/DsrC/DsvC family sulfur relay protein; MetB, cystathionine gamma-synthase family protein; MetX, homoserine O-acetyltransferase; PhsA, thiosulfate reductase; DmsB, anaerobic dimethyl sulfoxide reductase subunit B; SQR, sulfide: quinone oxidoreductase; SsuA, aliphatic sulfonate ABC transporter. More detailed information about proteins shown in this Figure was listed in the Supplementary Table 2.

To understand the underlying mechanisms of strain CS1 responding to different sulfur sources, we conducted a proteomic analysis of strain CS1 that cultured in the medium supplemented with Na_2_S_2_O_3_, Na_2_SO_3_ and Na_2_S, respectively. Since the growth status of *D*. *marinus* CS1 in different sulfur sources was inconsistent, we collected bacterial cells for proteome analysis when the OD_600_ value was about 0.08∼0.1 for both experimental and control groups. The proteomic results showed that compared to the control group total 1070, 1255 and 1012 proteins were significantly differentially expressed in the experimental groups containing Na_2_S_2_O_3_, Na_2_SO_3_ and Na_2_S (*P* < 0.05), respectively. Notably, most key enzymes associated with dissimilatory sulfate reduction (such as QmoA, QmoB, QmoC, AprA, AprB, Sat, DsrA and DsrB) were evidently down-regulated when cultured strain CS1 in the medium supplemented with Na_2_S_2_O_3_ (Fig. 1g), Na_2_SO_3_ (Fig. 1h) and Na_2_S (Fig. 1i), indicating that the supplement of Na_2_S_2_O_3_, Na_2_SO_3_ and Na_2_S suppressed the process of dissimilatory sulfate reduction in strain CS1. In contrast, the expressions of proteins related to thiosulfate and sulfide metabolisms, such as PhsA (thiosulfate reduction), MetB and SQR (sulfide oxidation), were significantly up-regulated when cultured strain CS1 in the medium supplemented with Na_2_S_2_O_3_ (Fig. 1g), Na_2_SO_3_ (Fig. 1h) and Na_2_S (Fig. 1i).

### *D. marinus* CS1 produces ZVS via metabolizing thiosulfate

It’s noting that some obvious white substances were observed in the medium supplemented with 40 mM Na_2_S_2_O_3_ when cultured *D. marinus* CS1 for about 20 days (Fig. 2a). In our previous study, we found a deep-sea bacterium *Erythrobacter flavus* 21-3 that isolated from the same sampling site of strain CS1 could oxidize Na_2_S_2_O_3_ to form ZVS through a novel sulfur oxidation pathway^17^. To clarify whether these white substances produced by strain CS1 were also ZVS, the SEM and EDS assays were conducted for initial assessment. SEM results showed that the white substances formed regular crystals (Fig. 2b), which were further identified as elemental sulfur by EDS (Fig. 2c). On the other hand, Raman spectrum showed that three strong peaks at 154, 221 and 475 cm^-1^ were identified toward the white substances produced by *D. marinus* CS1 (Fig. 2d). According to the cyclooctasulfur standard, these peaks corresponded to the bending and stretching modes of the 8-fold ring, and belonged to the typical characteristics of S_8_ (Fig. 2e)^6, 8, 17^. Meanwhile, the formation of ZVS mediated by strain CS1 was tracked across the whole two-month incubation period in the medium supplemented with thiosulfate. The results showed that ZVS could be detected after three-week incubation and its amount reached a stationary phase after five-week incubation (Fig. 3a), which presented a similar pattern with the growth curve of strain CS1 grown in the medium supplemented with thiosulfate (Fig. 1a). Accordingly, the concentration of thiosulfate decreased along with the formation of ZVS, while the concentration of sulfate almost remained unchanged (Fig. 3a). In comparison, strain CS1 could not form any ZVS in the medium absent of extra thiosulfate (Fig. 3b), while the concentration of sulfate decreased along with the bacterial growth (Fig. 3b). All the above results indicated that strain CS1 drove the formation of ZVS via metabolizing thiosulfate.

**Fig. 2.**
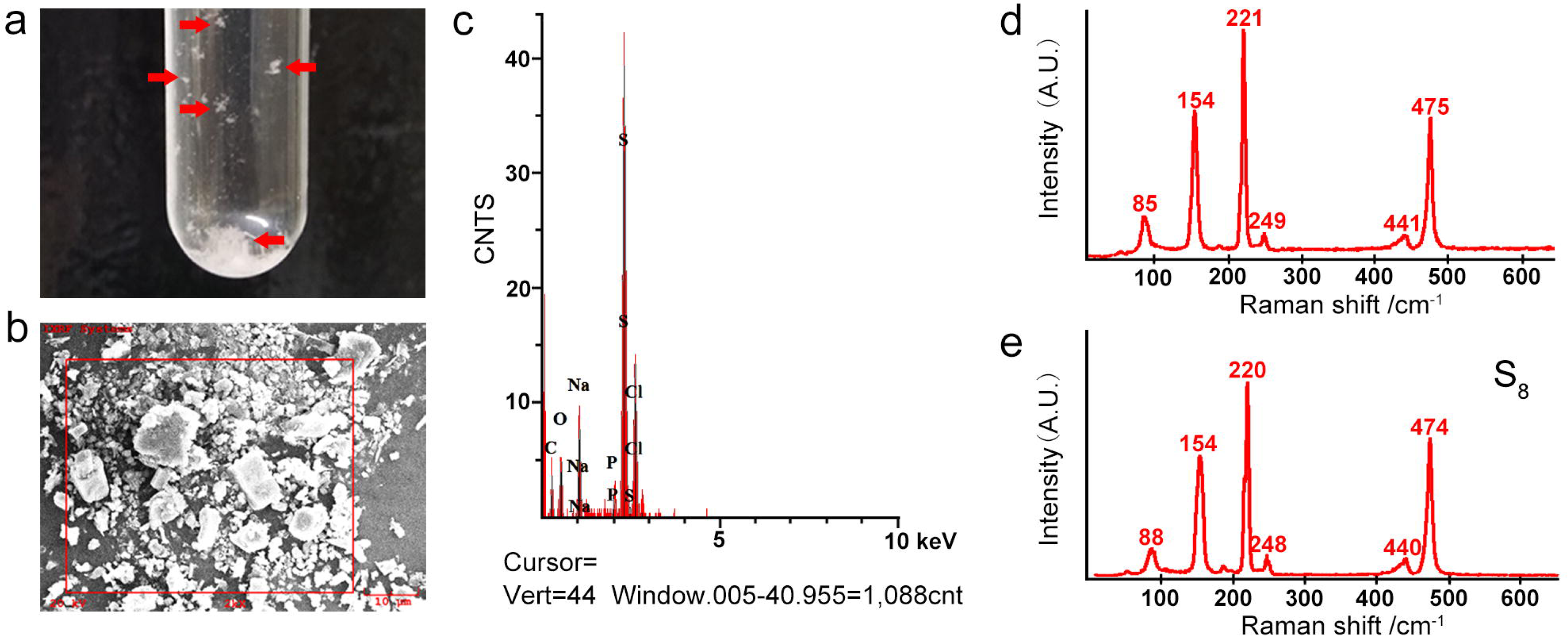
***D*. *marinus* CS1 produces ZVS when cultured in the medium supplemented with 40 mM Na_2_S_2_O_3_.** (a) Formation of obvious white substances by *D*. *marinus* CS1 when cultured in the medium supplemented with 40 mM Na_2_S_2_O_3_ (indicated with red arrows). (b) SEM observation of white substances produced by *D*. *marinus* CS1 shown in panel a. (c) Identification of major sulfur composition of white substances produced by *D*. *marinus* CS1 via energy dispersive spectrum (EDS) assay. (d) Confirmation of S_8_ configuration of white substances produced by *D*. *marinus* CS1 via Raman spectra assay. (e) Raman spectrum of standard S_8_.

**Fig. 3.**
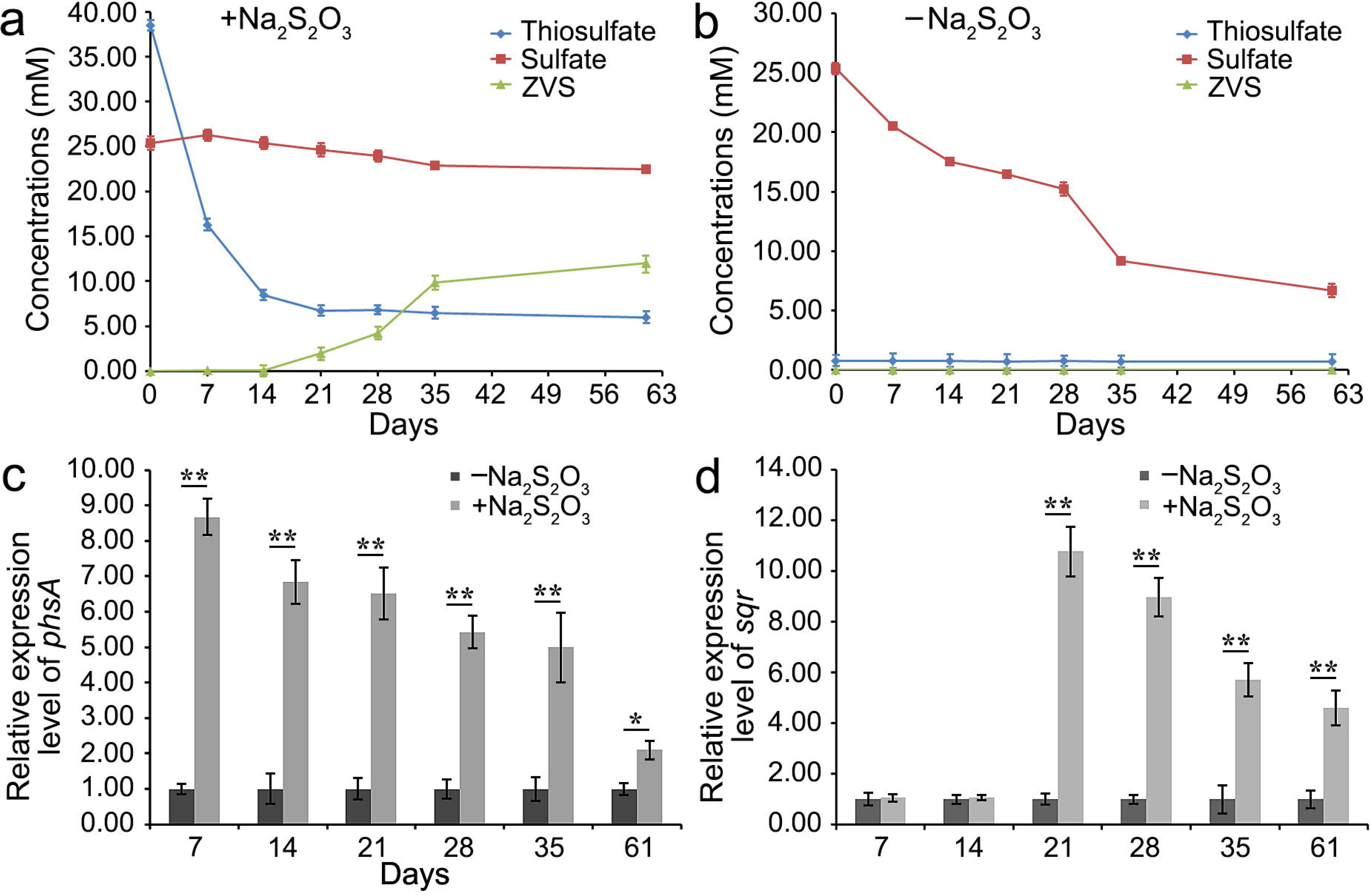
Monitoring the dynamics of concentrations of different sulfur intermediates and expression levels of *phsA* and *sqr* in the medium supplemented with 40 mM Na_2_S_2_O_3_ across the whole two-month incubation period. Dynamics of concentrations of sulfate, thiosulfate and ZVS in the medium supplemented with (a) or without (b) 40 mM Na_2_S_2_O_3_ across the whole two-month incubation period. The error bars indicate the standard deviation (S.D.) from three different biological replicates. Dynamics of the relative expression levels of *phsA* (c) and *sqr* (d) by qRT-PCR in the medium supplemented with or without 40 mM Na_2_S_2_O_3_ across the whole two-month incubation period. All data are relative to the expression levels found in the control group ± the standard error (N = 4). *, *P* < 0.05; **, *P* < 0.01.

### PhsA and SQR play key roles in driving ZVS formation in *D. marinus* CS1

Next, we sought to ask what determines the formation of ZVS in strain CS1 in the presence of thiosulfate. Given PhsA and SQR existing in strain CS1 and their evident expression up-regulation in the conditions supplemented with different sulfur sources (Figs. 1g-i), we thus speculate whether PhsA might metabolize thiosulfate to sulfide which in turn is oxidized to ZVS by SQR. To verify this assumption, we analyzed the dynamics of the expression levels of *phsA* and *sqr* along with the formation of ZVS as shown in Figure 4A. The results showed the expression level of *phsA* presented a decreasing trend from the beginning to the end of the incubation period though it was significantly up-regulated when compared to the control (Fig. 3c). On the other hand, the expression level of *sqr* was only markedly up-regulated after three-week incubation when compared to the control and then presented a decreasing trend till the end of the incubation period (Fig. 3d), which showed a similar pattern of ZVS formation as shown in Figure 4A. Indeed, the above results were consistent well with our speculation that PhsA and SQR respectively catalyze thiosulfate to form sulfide and then ZVS.

**Fig. 4.**
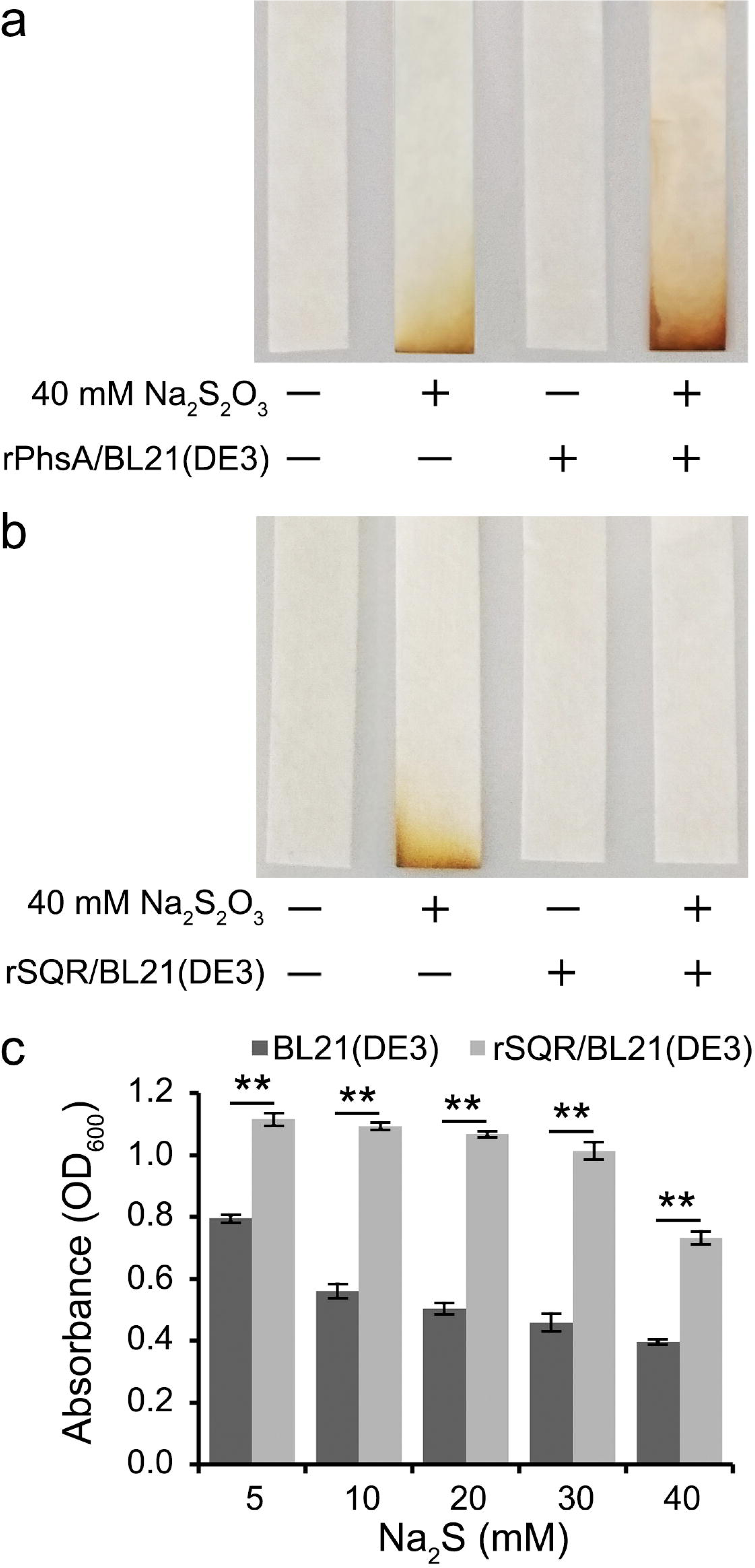
Functional assays of key proteins driving formation of ZVS in *D*. *marinus* CS1. (a) Overexpression of PhsA in *E*. *coli* promotes the transformation of S_2_O_3_^2-^ to H_2_S. *E*. *coli* cells without or with expression of PhsA were incubated in LB medium supplemented with 40 mM Na_2_S_2_O_3_ for 24 h. H_2_S accumulation was detected with lead-acetate paper strips. (b) Overexpression of SQR in *E*. *coli* promotes the transformation of H_2_S to other forms. *E*. *coli* cells without or with expression of SQR were incubated in LB medium supplemented with 40 mM Na_2_S_2_O_3_ for 24 h. H_2_S accumulation was detected with lead-acetate paper strips. (c) Growth assays of *E*. *coli* cells without or with expression of SQR in LB medium supplemented with respective 5 mM,10 mM, 20 mM, 30 mM and 40 mM Na_2_S for 24 h. ** means *P* < 0.01.

Given the absence of genetic operation system of strain CS1, we further verified the above functions of PhsA and SQR in *Escherichia coli*. First, we respectively overexpressed *phsA* (E8L03_06385) and *sqr* (E8L03_05425) of *D*. *marinus* CS1 in *E*. *coli* BL21(DE3) cells (Supplementary Fig. 3). PhsA has been proven to catalyze the decomposition of thiosulfate into sulfite and H_2_S^39–41^. Accordingly, the overexpression of PhsA in *E*. *coli* significantly promoted the production of H_2_S (Fig. 4a), indicating that PhsA of strain CS1 indeed functioned as an enzyme that catalyzing thiosulfate to H_2_S. On the other hand, SQR has potentials to oxidize sulfide to ZVS as shown in Figure 1D. As expected, the overexpression of SQR could efficiently remove the H_2_S produced by *E*. *coli* when cultured in the medium supplemented with 40 mM Na_2_S_2_O_3_ (Fig. 4b), which benefits the bacterial cells to alleviate the toxin effects of H_2_S. We thus further analyzed the activity of sulfide oxidation mediated by SQR in *E*. *coli* BL21(DE3) that cultured in the medium supplemented with different concentrations of Na_2_S (5 mM, 10 mM, 20 mM, 30 mM and 40 mM). If SQR could convert sulfide to ZVS, the toxicity of sulfide to *E*. *coli* cells would be significantly weakened. Indeed, the overexpression of SQR in *E*. *coli* BL21(DE3) could significantly promote (*P* < 0.01) bacterial growth when compared with the control group regardless of the concentrations of Na_2_S supplemented in the medium (Fig. 4c). Therefore, we propose that PhsA and SQR might drive ZVS formation in strain CS1 via respectively catalyzing thiosulfate and sulfide.

As SQR is a key enzyme catalyzing sulfide to form ZVS, we further analyzed SQR homologs identified in different microbes. In total, six types (Type I to Type VI) have been identified in bacteria, archaea and eukaryotes^19^. Based on the phylogenetic analysis, SQR in *D*. *marinus* CS1 was clustered into the branch of Type III SQRs with two conserved amino acid residues at Cys159 and Cys331 (Supplementary Fig. 4). Homologous sequences of SQR in strain CS1 were also identified in other SRB-D species that including *Desulfovibrio indonesiensis*, *Desulfohalovibrio alkalitolerans*, *Desulfatirhabdium butyrativorans*, *Desulfospira joergensenii*, *Pseudodesulfovibrio* sp. SRB007 and *Pseudodesulfovibrio* sp. zrk46 (Supplementary Fig. 4). It’s noting that *Pseudodesulfovibrio* sp. SRB007 and *Pseudodesulfovibrio* sp. zrk46 are two deep-sea SRB that isolated from the same sampling site as that of strain CS1.

### Proteomic analysis of sulfur metabolism of *D*. *marinus* CS1 in the deep sea

As shown above, *D*. *marinus* CS1 responded to different sulfur-containing compounds and formed ZVS in the laboratorial condition. Given that strain CS1 is a typical deep-sea sulfate reducing bacterium, it is necessary to explore its sulfur metabolisms in the deep-sea environment to mimic its lifestyle in the isolation niches. Taking the advantage of a cruise in May 2020, we cultured strain CS1 in the deep-sea cold seep for 10 days (Fig. 5a). Based on the environmental parameters of sites of *in situ* test and strain CS1 isolation (Supplementary Table S3), the two sites possessed pretty similar conditions. After 10 days incubation, bacterial cells in different groups were collected and performed proteomic assays after verification of their purity. As expected, according to the proteome data, the expressions of most key proteins associated with sulfate reduction (both assimilatory and dissimilatory) in the “In situ” group were significantly up-regulated compared to those in the laboratorial condition (Fig. 5b), strongly indicating the dominant function of sulfate reduction for strain CS1 to thrive in the deep-sea environment. Surprisingly, the expression of SQR was most up-regulated in the “In situ” group when compared to that cultivated in the laboratorial condition (Fig. 5b). Given that SQR was also significantly up-regulated when stimulating strain CS1 with thiosulfate (Fig. 3d) and sulfite (Fig. 1h) in laboratorial conditions and broadly distributed in different bacteria (Supplementary Figs. 4 and 5), we propose that SQR might play an essential role in driving sulfide oxidation in strain CS1 and other microbes. Meanwhile, the expression of PhsA was also evidently up-regulated in the “In situ” group (Fig. 5b), indicating thiosulfate metabolization is a major metabolic pathway for strain CS1 in the deep-sea environment. Given the fact that the expressions of SQR and PhsA were simultaneously up-regulated, we prefer the proposal that strain CS1 could form ZVS in the deep-sea environment.

**Fig. 5.**
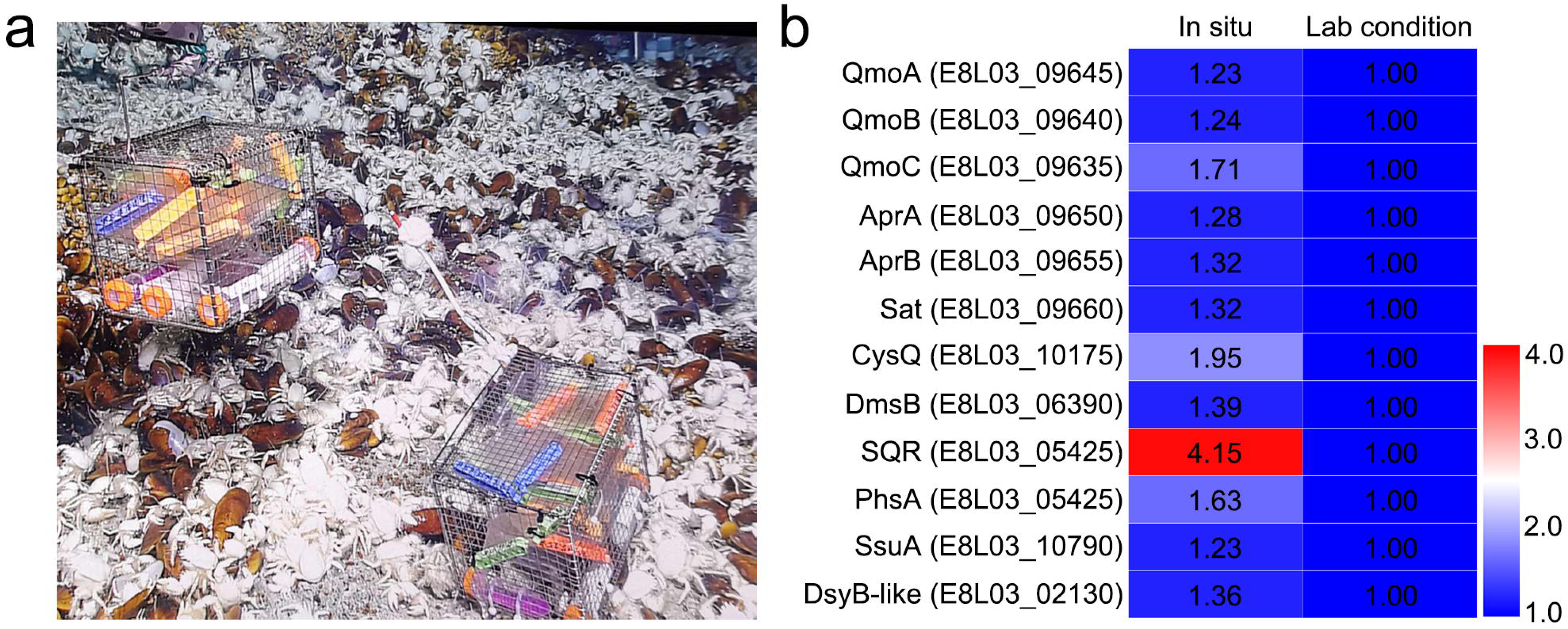
Proteomic analyses of sulfur metabolism of *D*. *marinus* CS1 that cultured in the deep-sea cold seep. (a) Representative picture showing the *in situ* experimental apparatus used in the deep-sea cold seep. (b) Proteome based heat map showing all different expressed proteins involved in sulfur metabolism after a 10-day incubation of *D*. *marinus* CS1 in the “In situ” group compared with “Lab condition” group. The numbers in the heat map represent the fold change of proteins compared to Lab conditions group. Abbreviations: DsyB-like, MTHB methyltransferase; dimethylsulphoniopropionate biosynthesis enzyme; CysQ, 3’(2’), 5’-bisphosphate nucleotidase. Other abbreviations are the same as shown in Figure 1. More detailed information about proteins shown in this Figure was listed in the Supplementary Table 2.

Overall, based on our present results, a proposed model towards central sulfur metabolisms of *D*. *marinus* CS1 was constructed (Fig. 6). First, sulfate is transported into the cells and then reduced to sulfite through both dissimilatory and assimilatory reduction pathways. Thereafter, sulfite is further reduced to sulfide mediated by the DSR complex via a typical sulfite dissimilatory reduction pathway. Meanwhile, thiosulfate is reduced to sulfide by PhsA. Finally, part of the generated sulfide is used for amino acids (e.g. cysteine and methionine) synthesis, and the rest is oxidized to polysulfide or even ZVS by SQR. The formed ZVS is finally exported to the outside of cells, contributing to form the mass ZVS around cold seep that observed in our previous reports^6, 17^.

**Fig. 6.**
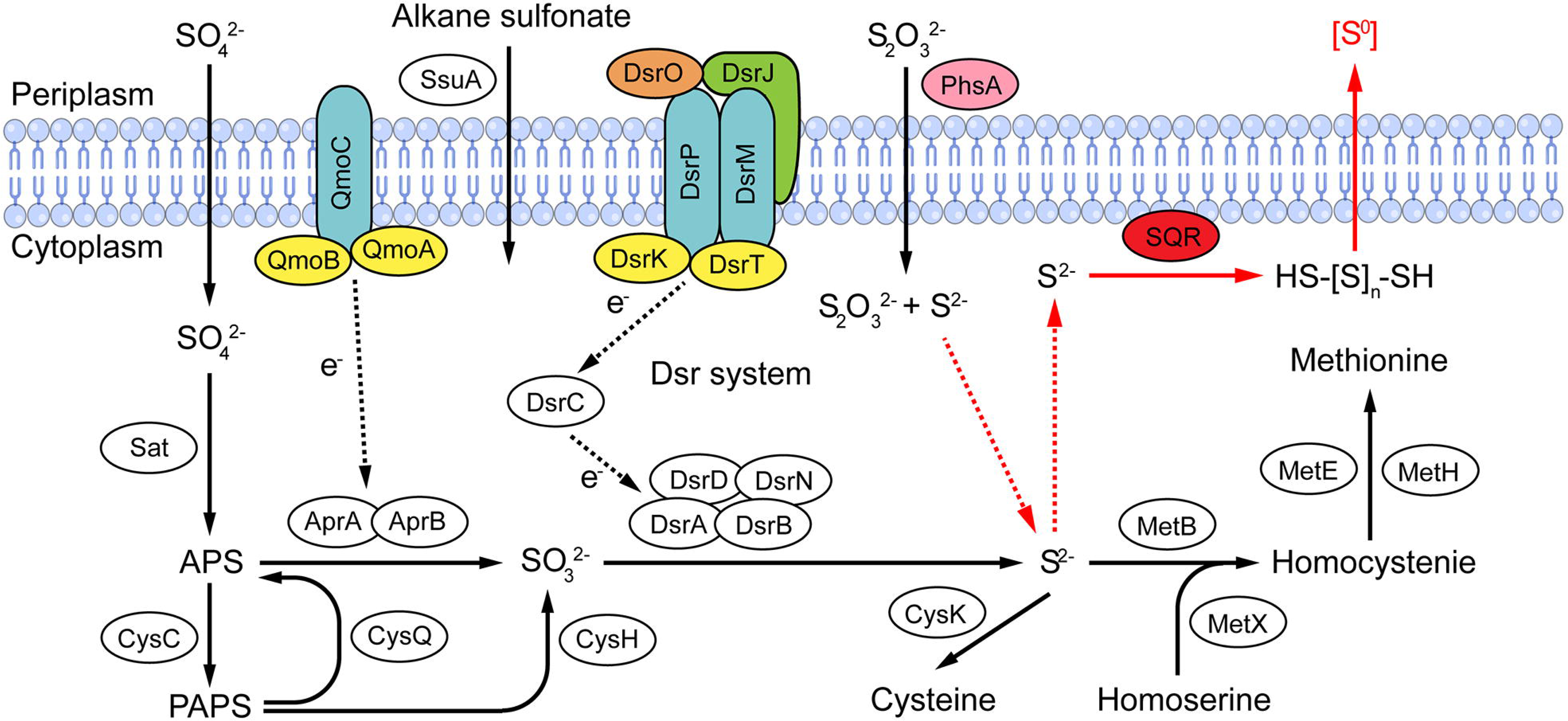
Proposed model related to sulfur metabolism and ZVS formation of *D*. *marinus* CS1. Black solid lines represent the typical sulfate reduction pathway present in *D*. *marinus* CS1. Black dashed lines represent the direction of electron transfer. Red lines represent the unique ZVS formation process present in *D*. *marinus* CS1. Abbreviations: APS, adenosine 5’-phosphosulfate; PAPS, 3’-phosphoadenosine-5’-phosphosulfate (3’-phosphoadenylylsulfate); CysC, adenylyl-sulfate kinase; CysH, phosphoadenosine phosphosulfate reductase family protein; DsrD, dissimilatory sulfite reductase-asociated protein; DsrK, [DsrC]-trisulfide reductase; DsrM, sulfate reduction electron transfer complex DsrMKJOP subunit; DsrN, cobyrinate a,c-diamide synthase; DsrJ, sulfate reduction electron transfer complex DsrMKJOP subunit; DsrO, 4Fe-4S dicluster domain-containing protein; DsrP, polysulfide reductase; DsrT, dissimilatory sulfite reductase system component; MetE, 5-methyltetrahydropteroyltriglutamate-homocysteine S-methyltransferase; MetH, methylenetetrahydrofolate reductase [NAD(P)H]; CysK, cysteine synthase. Other abbreviations are the same as shown in Figures 1 and 5. More detailed information about proteins shown in this figure was listed in the Supplementary Table 2.

### SRB potentially contribute to the formation of ZVS in deep sea

Based on our above results, *D*. *marinus* CS1 was demonstrated to form ZVS in the laboratorial conditions and possible deep-sea environment. We next sought to clarify the abundance of SRB in the deep-sea cold seep and their potentials for the formation of ZVS. As SRB belong to *D*-proteobacteria (SRB-D), the abundance of SRB-D was thus investigated by using the operational taxonomic units (OTUs) method with the sample collected from the surface (0-20 cm) of sediments. The results showed that the ration of SRB-D to the whole bacterial community was about 10% (Supplementary Fig. 6a). Among them, orders Desulfuromonadales and Desulfobacterales respectively accounted for 70% and 20%, while order Desulfovibrionales only accounted for less than 10% (Supplementary Fig. 6a). To obtain deeper insights into the distribution of genes associated with sulfur metabolisms in SRB-D, metagenomic sequencing was performed with samples collected from different depth intervals from the sedimental surface. As expected, genes associated SRB-D had a very high abundance in the cold seep sediments, whose percentages respectively accounted for 10.38%, 16.88%, 21.10%, 10.75% and 5.62% of the whole bacterial community in the samples C1, C4, C2, C3 and C5 (Supplementary Fig. 6b). After careful annotation and analyses of genes obtained from the metagenomic sequencing, we found that key genes responsible for both sulfate dissimilatory and assimilatory reduction pathways broadly distributed in the metagenomes of SRB-D and other bacteria in different samples (Supplementary Fig. 6c). And genes *sqr* and *phsA* could be identified in different samples with relative high proportions, strongly indicating the SRB-D in different depths of sediments have potentials to form ZVS.

## Discussion

Zero-valent sulfur (ZVS), in the form of elemental sulfur and dissolved polysulfide sulfur, is commonly measured in the highly reducing, sulfidic environments that characterize AOM ecosystems including the cold seeps^25^. SRB, a kind of important population inhabiting in cold seeps, have a pivotal role in the sulfur cycle, from which the generation of ZVS represents a novel pathway^24, 28^. ZVS generation from SRB was proposed to be mediated by the dissimilatory sulfate reduction under unfavorable conditions, e.g., inhibitive high-concentrations of sulfide^24, 28^. Hence, sulfide is a key intermediate for SRB to produce ZVS. It is noting that sulfide could be generated from thiosulfate via a reduction process catalyzed by PhsA, a kind of thiosulfate reductase (Supplementary Fig. 2d). Therefore, it is possible that SRB could also generate ZVS from thiosulfate in addition to sulfate. However, till to date, there is no any study showing the process and mechanisms of ZVS production from thiosulfate mediated by SRB. In the present study, we report for the first time that *D*. *marinus* CS1, a typical deep-sea sulfate reducing bacterium, could generate ZVS from thiosulfate coordinately mediated by PhsA and SQR. In this process, PhsA catalyzes thiosulfate to form sulfide, which is then oxidized by SQR to form ZVS.

Thiosulfate has been mentioned as an important shunt in marine environment for coupling of reductive and oxidative pathways of the sulfur cycle^32^. And the reduction of thiosulfate is a crucial process for anaerobic energy metabolism of SRB in marine sediments^42, 43^. In surface marine sediments (0∼10 cm depth), totally 15%∼50% of thiosulfate is reduced by sulfate reducing microorganisms, and approximately 30%∼60% of sulfide is produced during this process^32^. PhsA or its homologs are crucial for catalyzing thiosulfate to sulfide, which greatly contributes to an intraspecies sulfur cycle that drives S_0_ respiration in different bacteria^39, 44^. For *D*. *marinus* CS1, *phsA* was identified in its genome and was proposed to encode PhsA protein to reduce thiosulfate to sulfide (Supplementary Fig. 2d). Indeed, its expression level was significantly up-regulated in the medium supplemented with different sulfur sources in the laboratorial (Figs. 1g-1i) and deep-sea *in situ* (Fig. 5b) conditions, strongly indicating it is essential for sulfur cycling of *D*. *marinus* CS1. Especially, its expression was evidently up-regulated across the whole two-month incubation period in the presence of thiosulfate (Fig. 3c) and it was believed to play an indispensable role for strain CS1 to reduce thiosulfate to generate sulfide. Consistently, its mediation of thiosulfate to sulfide was verified in *E. coli* cells with the overexpression system (Fig. 4a).

As we known, sulfide is a highly toxic compound for microorganisms and eukaryotes^45, 46^. However, on the other hand, sulfide is also a very common intermediate of sulfur cycle in most microorganisms, and microbes have evolved different strategies to transform it to other forms given its strong toxicity. Indeed, the addition of Na_2_S significantly slowed down the growth of strain CS1 and it took a very long time and much energy for the bacterial cells to remove the toxicity effects of sulfide (Fig. 1a). SQR, a kind of oxidoreductase, enables to oxidize sulfide to ZVS and has potentials to alleviate the toxicity of sulfide^19^. Accordingly, we identified a gene encoding SQR in the genome of strain CS1 (Supplementary Fig. 2d). Notably, the expression dynamics of SQR showed a very similar pattern to the formation of ZVS when cultured strain CS1 in the medium supplemented with thiosulfate (Figs. 3a and 3d), indicating SQR is closely related to the formation of ZVS from thiosulfate in strain CS1. Based on the function of SQR, we believe that SQR identified in strain CS1 is capable of oxidize sulfide to ZVS for toxicity removal, which was confirmed by the effects of SQR overexpressed in *E*. *coli* BL21(DE3) to reduce the toxicity of Na_2_S from 5 mM to 40 mM (Fig. 4). It is noting that another member of *Desulfovibrio* genus (*D. pigers* Vib-7) could not grow in the medium containing 6 mM or higher concentration of sulfide^47^, while strain CS1 could tolerate up to 10 mM sulfide (Fig. 1a). Interestingly, the homologous sequence of SQR in strain CS1 was absent in the genome of *D*. *piger* (LT630450.1). Therefore, it is reasonable to deduce that SQR is an essential protein to drive sulfide oxidation to ZVS and thereby increasing the tolerance of *D*. *marinus* CS1 to sulfide.

Notably, in the deep-sea *in situ* environment, the expressions of both PhsA and SQR were markedly up-regulated (Fig. 5b). Due to the time limitation of our cruise, we were unable to culture *D*. *marinus* CS1 for a longer time (e.g. up to 60 days or longer) to observe the formation of ZVS. However, based on the fact that both expressions of PhsA and SQR were significantly up-regulated, we firmly believe that strain CS1 should form ZVS in the deep-sea cold seep. Through the metagenomic analysis, we found homologues of SQR of strain CS1 also broadly distributed in other SRB species and bacteria inhabiting in the same cold seep environment of the South China Sea (Supplementary Fig. 5, Supplementary Dataset 1). Moreover, homologs of key proteins (including PhsA and SQR) involved in sulfur cycle in *D*. *marinus* CS1 were found to widely exist in the metagenomes of SRB and other bacteria that living in the sediments of the South China Sea (Supplementary Fig. 6c). Therefore, we propose that the novel ZVS formation pathway from thiosulfate metabolization mediated by PhsA and SQR is used by a lot of SRB or even other bacteria in the deep-sea environments, which might play an undocumented role in the sulfur cycle and encourages the re-evaluation of the contribution of SRB to the formation of ZVS in the deep ocean.

## Methods

### Isolation and cultivation of *D. marinus* CS1

To isolate SRB from the deep-sea environment, cold seep sediment samples were collected by R/V *KEXUE* in the South China Sea (119°17’04.956’’E, 22°06’58.384’’N) at a depth of approximately 1,143 m in September 2017 (Supplementary Table S3). The samples were cultured by using the modified sulfate reducing medium (SRM) that containing 6.5 g PIPES (C_8_H_18_N_2_O_6_S_2_), 2.7 g MgSO_4_·7H_2_O, 4.3 g MgCl_2_·6H_2_O, 0.25 g NH_4_Cl, 0.5 g KCl, 0.14 g CaCl_2_, 0.14 g K_2_HPO_4_·3H_2_O, 0.01 g Fe(NH_4_)_2_(SO_4_)_2_·6H_2_O, 0.1 g CH_3_COONa, 2.24 g CH_3_CHOHCOONa, 20 mM absolute ethanol, 1 mL trace elements solution (Supplementary Table S4), 1 mL vitamins solution (Supplementary Table S5), 0.5 g cystine and 0.001 g resazurin in 1 liter filtered sea water, and 15 g/L agar was added to prepare the corresponding solid medium. After a month anaerobic enrichment at standard atmospheric pressure, a 50 µL culture was spread on the solid SRM medium prepared in the Hungate tubes, which were further anaerobically incubated at 28 °C for 7 days. Individual colonies were picked respectively by sterilized bamboo sticks and then cultured in the SRM broth. Strain CS1 was isolated and purified by the Hungate roll-tube method for several rounds until considered to be axenic. Genomic DNA extraction and PCR amplification of the 16S rRNA gene sequence of strain CS1 were performed as previously described previously^17^.

### Electron microscopic analysis

The morphological characteristics of *D*. *marinus* CS1 were observed by scanning electron microscope (SEM) (S-3400N; Hitachi, Japan) and transmission electron microscope (TEM) (HT7700; Hitachi, Japan). The ZVS produced by strain CS1 in the medium supplemented with Na_2_S_2_O_3_ was identified via Energy-Dispersive Spectrum (EDS) (model 550i, IXRF systems, USA) equipment with SEM and Raman spectra confocal microscope (WITec alpha300 R system; WITec Company, Germany), respectively, as described in our previous study^17^. After incubated in the medium supplemented with 40 mM Na_2_S_2_O_3_ for 30 days, the milky white supernatant in strain CS1 medium was collected by centrifugation (5,000 *g*, 10 min) and lyophilized, then the pellet was used for EDS analysis at an accelerating voltage of 5 keV for 30 s and Raman spectra.

### Genome sequencing and annotation

Genomic DNA was extracted from *D. marinus* CS1 that cultured for 7 days at 28 °C. The harvested DNA was detected by the agarose gel electrophoresis and quantified by Qubit 3.0 (Thermo Fisher Scientific, USA). Whole-genome sequence determinations of strain CS1 were carried out with the PacBio (Pacific Biosciences, USA) and Illumina MiSeq (Illumina, USA) sequencing platform. The genome of strain CS1 was sequenced by PacBio platform (PacBio, USA) using single molecule real-time (SMRT) technology. Sequencing was performed at the Beijing Novogene Bioinformatics Technology Co., Ltd. The low quality reads were filtered by the SMRT Link v5.0.1 and the filtered reads were assembled to generate one contig without gaps and was manually circularized by deleting an overlapping end.

The genome relatedness values were calculated by Average Nucleotide Identity (ANI) based on BLASTN algorithm with recommended species criterion cut-offs 95% (JSpecies WS, http://jspecies.ribohost.com/jspeciesws/) and the amino acid identity (AAI) based on AAI-profiler with values above 95∼97 % correspond to the same species (http://ekhidna2.biocenter.helsinki.fi/AAI/). To determine the phylogenetic position of *D*. *marinus* CS1, the 16S rRNA gene sequence was analyzed by the BLAST programs (https://blast.ncbi.nlm.nih.gov/Blast.cgi), and the phylogenetic tree was reconstructed with MEGA X^48^.

### Growth assays of *D*. *marinus* CS1 in the medium supplemented with different sulfur sources

Growth assays were performed under the standard atmospheric pressure. Briefly, 15 mL fresh *D*. *marinus* CS1 was cultured in SRM supplemented with or without 20 mM Na_2_SO_4_, 40 mM Na_2_S_2_O_3_, 10 mM Na_2_SO_3_ and 10 mM Na_2_S for two months at 28 °C in 2 L anaerobic bottles, respectively. Each condition had three replicates. Bacterial growth status was monitored by measuring the absorbance value at 600 nm (OD_600_). For the morphological observation, cells of strain CS1 with OD_600_ values at 0.08∼0.1 were collected and recorded under the TEM as described above.

### Proteomic analysis

Proteome analysis was performed by PTM BIO (PTM Biolabs Inc., China). Briefly, cell suspensions of *D*. *marinus* CS1 with an OD_600_ value of 0.08∼0.1 were collected from different groups at different time points: 7 days for control group and Na_2_SO_4_ supplement group, 14 days for Na_2_S_2_O_3_ supplement group and Na_2_S supplement group, and 42 days for Na_2_SO_3_ supplement group, respectively. Thereafter, the cells were checked by 16S rRNA sequencing to confirm the purity of the culture and performed further proteomic analysis. For proteome analyses, the cells of strain CS1 were washed with 10 mM phosphate buffer solution (PBS pH 7.4), resuspended in lysis buffer (8 M urea, 1% Protease Inhibitor Cocktail) and disrupted by sonication. The remaining debris was removed by centrifugation at 12,000 *g* at 4 °C for 10 min. Finally, the supernatant was collected and the protein concentration was determined with a BCA protein assay kit (Thermo Fisher Scientific, USA) according to the manufacturer’s instructions. Trypsin digestion, TMT labeling, HPLC fractionation, LC-MS/MS analysis, database search and bioinformatics analysis are detailedly described in the supplementary information. Analysis of the differentially expressed proteins was performed using HemI software^49^.

To perform the proteomic analysis with the cells that cultured in the deep-sea cold seep, strain CS1 was firstly cultured in SRM for 7 days under laboratorial conditions, and then was separated into two parts: one was equally divided into three dialysis bags (8,000-14,000 Da cutoff, which allowing the exchanges of substances smaller than 8,000 Da but preventing bacterial cells from entering or leaving the bag; Solarbio, China) as the “In situ” group; the other part was equally divided into three anaerobic bottles and incubated for 10 days in the laboratorial conditions as the control group. The “In situ” group was placed simultaneously in the cold seep (E119°17′04.429, N22°07′01.523″) for 10 days in the June 2020 during the cruise of R/V *KEXUE*. After 10 days incubation in the deep sea, the bags were taken out and the cells were immediately collected and saved in the −80 °C freezer. 16S rRNA sequencing was performed to ensure the purity of cells of strain CS1 that collected from the *in situ* cultivation. The proteomic assays were performed as described above.

### Analytical techniques for the determination of different sulfur compounds and quantitative real-time PCR (qRT-PCR) assay

To detect the changes of concentrations of thiosulfate and sulfate, 50 mL cultures of *D*. *marinus* CS1 was collected from groups supplemented with or without 40 mM Na_2_S_2_O_3_ at the 7^th^, 14^th^, 21^st^, 28^th^, 35^th^ and 61^st^ day, respectively. Meanwhile, the growth status was monitored. After centrifugation at 12,000 *g* at 4 °C for 30 min, concentrations of thiosulfate and sulfate in the supernatant were respectively measured by iodometric and barium sulfate turbidimetry as described previously^50, 51^. The concentration of ZVS (S_8_) in the medium was detected according to the method described previously^52^. Briefly, 1 mL cultured medium was extracted three times using a total of 5 mL chloroform. The extracted sample was measured on a UV-Vis spectrometer (Infinite M1000 Pro; Tecan, Männedorf, Switzerland) at 270 nm^52^.

To perform the qRT-PCR assay, total RNAs of cells collected at different time points were extracted with TRIzol reagent (Invitrogen, USA). First-strand cDNA synthesis was carried out with ReverTra Ace® qPCR RT Master Mix (TOYOBO, Japan) based on the manufacturer’s instructions. The expression levels of *sqr* and *phsA* were determined using qRT-PCR on different cDNA samples obtained from cultures as described above. Specific primers were designed according to the corresponding sequences in the genome of *D*. *marinus* CS1 (Supplementary Table S6). The comparative threshold cycle (CT) (2^-ΔΔ^) method was used to analyze the expression level^53^. Two 16S rRNA gene primers for *D*. *marinus* CS1 (Supplementary Table S6) were used as internal controls to verify successful transcription and to calibrate the cDNA template for corresponding samples. qRT-PCR was performed using a Quant Studio^TM^ 6 Flex (Life Technologies, USA), and the collected data were analyzed with the system’s accompanying SDS software. Dissociation curve analysis of amplification products was performed at the end of each PCR to confirm that only one PCR product was amplified and detected. All data were given in terms of relative mRNA expressed as means ± standard error (N=4).

### Functional assays of PhsA and SQR of *D*. *marinus* CS1

To detect the functions of SQR and PhsA that identified in *D*. *marinus* CS1, the genes encoding these two proteins were respectively cloned and overexpressed in *E*. *coli*. First, the open reading frame of *sqr* or *phsA* was amplified from the *D*. *marinus* CS1 genome using the KOD One ^TM^ PCR Master Mix (TOYOBO, Japan) with corresponding primers (Supplementary Table S6). The PCR product was purified by using a DNA Gel Extraction Kit (TsingKe, China), and then was cloned in the plasmid pMD19-T simple (TAKARA, Japan). The DNA fragment was digested with *Eco*RI/*Xho*I and *Bam*HI/*Eco*RI (Thermo Fisher Scientific, USA), respectively, and ligated into the same restriction enzymes sites of expression vector pET28a (+) (Merck, Germany). The recombinant plasmids were transformed into competent cell *E*. *coli* BL21(DE3) (TsingKe, China), and transformants were incubated in Luria-Bertani broth (10 g NaCl, 10 g tryptone and 5 g yeast extract per liter of Milli-Q water) supplemented with 50 μg/mL kanamycin at 37 °C. Protein expression was induced at an OD_600_ around 0.6 with 0.1 mM isopropyl-1-thio-β-D-galactopyranoside (ITPG), and the cells were cultured for further 20 h at 16 °C. The resultant proteins were separated by SDS-PAGE, and visualized with Coomassie Bright Blue R250 staining.

Different concentrations of Na_2_S (5 mM, 10 mM, 20 mM, 30 mM and 40 mM) was respectively added in LB medium inoculated with *E*. *coli* BL21(DE3) containing the recombinant plasmid *sqr/*pET28a (+). The *E*. *coli* BL21(DE3) cells transformed with plasmid pET28a (+) were used as the negative control. After induced with 0.1 mM IPTG, *E*. *coli* BL21(DE3) cells overexpressing with or without recombinant SQR (rSQR) were cultured at 37 °C for 24 h. And then the growth statuses of *E*. *coli sqr/*pET28a (+)/BL21(DE3) (overexpressing rSQR) and the negative control *E*. *coli* pET28a (+)/BL21(DE3) that treated with different concentrations of Na_2_S, were measured on the Infinite M1000 Pro UV-Vis spectrometer (Tecan, Switzerland) at 600 nm (OD_600_). The lead acetate test papers were used to detect the amount of hydrogen sulfide (H_2_S) which reflecting the capability of PhsA for reducing thiosulfate to producing H_2_S. *E*. *coli sqr/*pET28a (+)/BL21(DE3), *phsA/*pET28a (+)/BL21(DE3) and the negative control *E*. *coli* pET28a (+)/BL21(DE3) were cultured in the medium supplemented with 40 mM Na_2_S_2_O_3_, respectively. After induced with 0.1 mM IPTG, the production of H_2_S was detected after incubation at 37 °C for 24 h.

### Data availability

The genomic data of *D. marinus* CS1 have been deposited to the NCBI with the accession number of CP039543.1. The mass spectrometry proteomics data have been deposited to the Proteome Xchange Consortium via the PRIDE^54^ partner repository with the dataset identifier PXD023247. The raw metagenomic sequencing data have been deposited to NCBI Short Read Archive (accession numbers: SRR13052532, SRR13063401, SRR13336710, SRR13065122 and SRR13065132). The raw amplicon sequencing data have also been deposited to NCBI Short Read Archive (accession number: SRR13360429).

### Statistical analysis

The significant differences among groups were subjected to one-way analysis of variance (one-way ANOVA) and multiple comparisons by using the SPSS 18.0 program. A statistical significance was defined in our study by *P* < 0.05 (indicated by *\** in all figures) or *P* < 0.01 (indicated by ** in all figures).

## Acknowledgements

This work was funded by the Strategic Priority Research Program of the Chinese Academy of Sciences (Grant No. XDA22050301), China Ocean Mineral Resources R&D Association Grant (Grant No. DY135-B2-14), Key Deployment Projects of Center of Ocean Mega-Science of the Chinese Academy of Sciences (Grant No. COMS2020Q04), Major Research Plan of the National Natural Science Foundation (Grant No. 92051107), National Key R and D Program of China (Grant No. 2018YFC0310800), the Taishan Young Scholar Program of Shandong Province (tsqn20161051), and Qingdao Innovation Leadership Program (Grant No. 18-1-2-7-zhc) for Chaomin Sun. This study is also funded by the CAS Interdisciplinary Innovation Team (Grant No. JCTD-2018-12) and Open Research Project of National Major Science & Technology Infrastructure (*RV KEXUE*) (Grant No. NMSTI-KEXUE2017K01).

## Author contributions

RL and CS conceived and designed the study; RL conducted most of the experiments; YS helped to perform the *in situ* experiments; SX and XZ helped to perform the Raman spectra analysis; RL and CS lead the writing of the manuscript; all authors contributed to and reviewed the manuscript.

## References

1. Schippers, A. & Jorgensen, B. B. Biogeochemistry of pyrite and iron sulfide oxidation in marine sediments. Geochim Cosmochim Ac 66, 85–92 (2002).

2. Holmkvist, L., Ferdelman, T. G. & Jorgensen, B. B. A cryptic sulfur cycle driven by iron in the methane zone of marine sediment (Aarhus Bay, Denmark). Geochim Cosmochim Ac 75, 3581–3599 (2011).

3. Wasmund, K., Mussmann, M. & Loy, A. The life sulfuric: microbial ecology of sulfur cycling in marine sediments. Env Microbiol Rep 9, 323–344 (2017).

4. White, S. N. Laser Raman spectroscopy as a technique for identification of seafloor hydrothermal and cold seep minerals. Chem Geol 259, 240–252 (2009).

5. White, S. N., Dunk, R. M., Peltzer, E. T., Freeman, J. J. & Brewer, P. G. In situ Raman analyses of deep-sea hydrothermal and cold seep systems (Gorda Ridge and Hydrate Ridge). Geochem Geophy Geosy 7 (2006).

6. Zhang, X. et al. Development of a new deep-sea hybrid Raman insertion probe and its application to the geochemistry of hydrothermal vent and cold seep fluids. Deep-Sea Res Pt I 123, 1–12 (2017).

7. Lichtschlag, A., Kamyshny, A., Ferdelman, T. G. & deBeer, D. Intermediate sulfur oxidation state compounds in the euxinic surface sediments of the Dvurechenskii mud volcano (Black Sea). Geochim Cosmochim Ac 105, 130–145 (2013).

8. Berg, J. S., Schwedt, A., Kreutzmann, A. C., Kuypers, M. M. M. & Milucka, J. Polysulfides as intermediates in the oxidation of sulfide to sulfate by *Beggiatoa* spp. Appl Environ Microb 80, 629–636 (2014).

9. Gleeson, D. F. et al. Biosignature detection at an arctic analog to Europa. Astrobiology 12, 135–150 (2012).

10. Hamilton, T. L., Jones, D. S., Schaperdoth, I. & Macalady, J. L. Metagenomic insights into S(0) precipitation in a terrestrial subsurface lithoautotrophic ecosystem. Front Microbiol 5 (2015).

11. Ghosh, W. & Dam, B. Biochemistry and molecular biology of lithotrophic sulfur oxidation by taxonomically and ecologically diverse bacteria and archaea. FEMS Microbiol Rev 33, 999–1043 (2009).

12. Duzs, A., et al. A novel enzyme of type VI sulfide: quinone oxidoreductases in purple sulfur photosynthetic bacteria. Appl Microbiol Biot 102, 5133–5147 (2018).

13. Meyer, B., Imhoff, J. F. & Kuever, J. Molecular analysis of the distribution and phylogeny of the soxB gene among sulfur-oxidizing bacteria - evolution of the Sox sulfur oxidation enzyme system. Environ Microbiol 9, 2957–2977 (2007).

14. Sakurai, H., Ogawa, T., Shiga, M. & Inoue, K. Inorganic sulfur oxidizing system in green sulfur bacteria. Photosynthesis research 104, 163–176 (2010).

15. Dam, B., Mandal, S., Ghosh, W., Das Gupta, S. K. & Roy, P. The S4-intermediate pathway for the oxidation of thiosulfate by the chemolithoautotroph *Tetrathiobacter kashmirensis* and inhibition of tetrathionate oxidation by sulfite. Research in microbiology 158, 330–338 (2007).

16. Hensen, D., Sperling, D., Truper, H. G., Brune, D. C. & Dahl, C. Thiosulphate oxidation in the phototrophic sulphur bacterium *Allochromatium vinosum*. Molecular microbiology 62, 794–810 (2006).

17. Zhang, J. et al. A novel bacterial thiosulfate oxidation pathway provides a new clue about the formation of zero-valent sulfur in deep sea. ISME J 14, 2261–2274 (2020).

18. Lahme, S. et al. Comparison of sulfide-oxidizing *Sulfurimonas* strains reveals a new mode of thiosulfate formation in subsurface environments. Environ Microbiol 22, 1784–1800 (2020).

19. Marcia M, Ermler U, Peng GH & Michel H. A new structure-based classification of sulfide: quinone oxidoreductases. Proteins 78, 1073–1083 (2010).

20. Lencina, A. M., Ding, Z. Q., Schurig-Briccio, L. A. & Gennis, R. B. Characterization of the Type III sulfide: quinone oxidoreductase from *Caldivirga maquilingensis* and its membrane binding. Biochim Biophys Acta 1827, 266–275 (2013).

21. Brito, J. A. et al. Structural and functional insights into sulfide: quinone oxidoreductase. Biochemistry 48, 5613–5622 (2009).

22. Muller, H., Marozava, S., Probst, A. J. & Meckenstock, R. U. Groundwater cable bacteria conserve energy by sulfur disproportionation. ISME J 14, 623–634 (2020).

23. van Vliet, D. M. et al. The bacterial sulfur cycle in expanding dysoxic and euxinic marine waters. Environ Microbiol (2020).

24. Fang, W. W., Liang, Z. W., Liu, Y. L., Liao, J. Y. & Wang, S. Q. Metagenomic insights into production of zero valent sulfur from dissimilatory sulfate reduction in a methanogenic bioreactor. Bioresour Technol Rep 383 (2019).

25. Milucka, J. et al. Zero-valent sulphur is a key intermediate in marine methane oxidation. Nature 491, 541–546 (2012).

26. Labrado, A. L., Brunner, B., Bernasconi, S. M. & Peckmann, J. Formation of large native sulfur deposits does not require molecular oxygen. Front Microbiol 10 (2019).

27. Yu, H. et al. Comparative genomics and proteomic analysis of assimilatory sulfate reduction pathways in anaerobic methanotrophic archaea. Front Microbiol 9 (2018).

28. Fang, W., Gu, M., Liang, D., Chen, G. H. & Wang, S. Generation of zero valent sulfur from dissimilatory sulfate reduction under methanogenic conditions. J Hazard Mater 383 (2020).

29. Perez-Lopez, R. et al. Sulfate reduction processes in salt marshes affected by phosphogypsum: Geochemical influences on contaminant mobility. J Hazard Mater 350, 154–161 (2018).

30. Grabarczyk, D. B. et al. Structural basis for specificity and promiscuity in a carrier protein/enzyme system from the sulfur cycle. Proc Natl Acad Sci U S A 112, E7166–7175 (2015).

31. Anantharaman, K. et al. Expanded diversity of microbial groups that shape the dissimilatory sulfur cycle. ISME J 12, 1715–1728 (2018).

32. Jorgensen, B. B. A thiosulfate shunt in the sulfur cycle of marine sediments. Science 249, 152–154 (1990).

33. Feng, D. et al. Cold seep systems in the South China Sea: An overview. J Asian Earth Sci 168, 3–16 (2018).

34. Wegener, G., Krukenberg, V., Riedel, D., Tegetmeyer, H. E. & Boetius, A. Intercellular wiring enables electron transfer between methanotrophic archaea and bacteria. Nature 526, 587–590 (2015).

35. McGlynn, S. E., Chadwick, G. L., Kempes, C. P. & Orphan, V. J. Single cell activity reveals direct electron transfer in methanotrophic consortia. Nature 526, 531–535 (2015).

36. Wu, Y., Qiu, J. W., Qian, P. Y. & Wang, Y. The vertical distribution of prokaryotes in the surface sediment of Jiaolong cold seep at the northern South China Sea. Extremophiles 22, 499–510 (2018).

37. Richter, M., Rossello-Mora, R., Glockner, F. O. & Peplies, J. JSpeciesWS: a web server for prokaryotic species circumscription based on pairwise genome comparison. Bioinformatics 32, 929–931 (2016).

38. Medlar, A. J., Toronen, P. & Holm, L. AAI-profiler: fast proteome-wide exploratory analysis reveals taxonomic identity, misclassification and contamination. Nucleic Acids Res 46, W479–W485 (2018).

39. Burns, J. L. & DiChristina, T. J. Anaerobic respiration of elemental sulfur and thiosulfate by *Shewanella oneidensis* MR-1 Requires psrA, a homolog of the phsA gene of *Salmonella enterica* serovar Typhimurium LT2. Appl Environ Microb 75, 5209–5217 (2009).

40. Aketagawa, J., Kobayashi, K. & Ishimoto, M. Purification and properties of thiosulfate reductase from *Desulfovibrio vulgaris*, Miyazaki F. J Biochem 97, 1025–1032 (1985).

41. Chen, Z. G. et al. The complete pathway for thiosulfate utilization in *Saccharomyces cerevisiae*. Appl Environ Microb 84 (2018).

42. Bak, F. & Cypionka, H. A novel type of energy metabolism involving fermentation of inorganic sulphur compounds. Nature 326, 891–892 (1987).

43. Jorgensen, B. B. & Bak, F. Pathways and microbiology of thiosulfate transformations and sulfate reduction in a marine sediment (kattegat, denmark). Appl Environ Microbiol 57, 847–856 (1991).

44. Wells, M. et al. Respiratory selenite reductase from B*acillus selenitireducens* strain MLS10. J Bacteriol 201 (2019).

45. Beauchamp, R. O., Jr., Bus, J. S., Popp, J. A., Boreiko, C. J. & Andjelkovich, D. A. A critical review of the literature on hydrogen sulfide toxicity. Crit Rev Toxicol 13, 25–97 (1984).

46. Blachier, F. et al. Luminal sulfide and large intestine mucosa: friend or foe? Amino acids 39, 335–347 (2010).

47. Kushkevych, I., Dordevic, D. & Vitezova, M. Toxicity of hydrogen sulfide toward sulfate-reducing bacteria *Desulfovibrio piger* Vib-7. Arch Microbiol 201, 389–397 (2019).

48. Kumar, S., Stecher, G., Li, M., Knyaz, C. & Tamura, K. MEGA X: Molecular evolutionary genetics analysis across computing platforms. Mol Biol Evol 35, 1547–1549 (2018).

49. Deng, W. K., Wang, Y. B., Liu, Z. X., Cheng, H. & Xue, Y. HemI: A toolkit for illustrating heatmaps. Plos One 9 (2014).

50. Trueper, H. G. & Schlegel, H. G. Sulphur metabolism in Thiorhodaceae. I. quantitative measurements on growing cells of Chromatium Okenii. Antonie van Leeuwenhoek 30, 225–238 (1964).

51. 51. Kelly, D. P. & Wood, A. P. in *Methods in Enzymology* Vol. 243 475-501 (Academic Press, 1994).

52. Houghton, J. L. et al. Thiosulfate oxidation by *Thiomicrospira thermophila*: metabolic flexibility in response to ambient geochemistry. Environ Microbiol 18, 3057–3072 (2016).

53. Livak, K. J. & Schmittgen, T. D. Analysis of relative gene expression data using real-time quantitative PCR and the 2(T)(-Delta Delta C) method. Methods 25, 402–408 (2001).

54. Perez-Riverol, Y. et al. The PRIDE database and related tools and resources in 2019: improving support for quantification data. Nucleic Acids Res 47, D442–D450 (2019).

