## Supplemental Figures and Tables for "A deep-sea sulfate reducing bacterium directs the formation of zero-valent sulfur via sulfide oxidation"

**Supplementary methods**

**Phylogenetic analysis**

To assess the type of SQR in *D*. *marinus* CS1, amino acid sequences of SQR homologs^1^^,^^2^ were extracted from NCBI databases (including Type I to VI SQRs) and our metagenomic sequencing data. After sequence alignment, a maximum-likelihood phylogenetic tree was constructed with the LG+G4+F model (-bb 1,000) using IQ-TREE. The sequence alignments for all trees were calculated using the MEGA X soft with Clustal W/MUSCLE program^3^. All trees were visualized using iTOL (v5)^4^.

**Metagenomic sequencing and amplicon analysis**

For metagenomics analysis, samples were collected from cold seep sediments at a depth of 1,146 m in the South China Sea during the cruise of the R/V *Kexue* in the July of 2018 (Supplementary Table S5). Samples were collected at depth intervals of 0-20 cm (sample C1), 20-40 cm (sample C4), 40-60 cm (sample C2), 120-140 cm (sample C3) and deeper than 280 cm (sample C5). Among them, sample C1 was collected by the Discovery remotely operated vehicle (ROV), sample C4 was collected by the television grab, and samples C2, C3 and C5 were collected by the gravity sampler. Concentrations of sulfate and CH_4_ in surface sediments of sampling sites were respective 28 mM and 2,642 μM that measured by Raman spectra and Hydro®CH4 (CONTROS, Norway) sensors. All sedimentary samples were dehydrated in an oven at 80 °C until completely dry. After grinding, sample powder was filtered through a 200-mesh screen. The filtrate was analyzed for chemical content of sulfur using an S8 Tiger X-ray fluorescence spectrometer (BRUKER, Germany). These five cold seep sediment samples (C1, C4, C2, C3 and C5, 20 g each) were further used for metagenomic analysis in BGI (BGI, China). Briefly, total DNAs from these samples were extracted using the DNeasy® PowerSoil® Pro Kit (Qiagen, Germany) and the integrity of DNA was evaluated by gel electrophoresis. 0.5 μg of each sample was used to prepare libraries. DNA was sheared into fragments between 50-800 bp in length using a Covaris E220 ultrasonicator (Covaris, UK); fragments between 150 bp and 250 bp were secreted using AMPure XP beads (Agencourt, USA) and repaired using T4 DNA polymerase (ENZYMATICS, USA). DNA fragments were ligated at both ends to T-tailed adapters and amplified for eight cycles. Finally, the amplification products were used to produce single-stranded, circular DNA libraries. All NGS (Next Generation Sequencing) libraries were sequenced on a BGISEQ-500 platform (BGI, China) to obtain 100 bp paired-end raw reads. Quality control was performed by SOAPnuke (v1.5.6) (setting: -l 20 -q 0.2 -n 0.05 -Q 2 -d -c 0 -5 0 -7 1)^5^. Gene prediction for metagenomics data was performed using Glimmer (v 3.02)^6^. The KEGG (Kyoto Encyclopedia of Genes and Genomes, Release 87.0), NR (Non-Redundant Protein Database databases, 20180814), Swiss-Prot (release-2017_07) and EggNOG (2015-10_4.5v) databases were used to annotate protein functions by default, and the best hits were chosen.

To understand the abundance of sulfate reducing bacteria belonging to *D*-proteobacteria in deep-sea cold seep, we selected the surface sediment sample C1 for operational taxonomic units (OTUs) sequencing performed by Genesky (Genesky Biotechnologies Inc., China). Total DNAs from sample C1 were extracted by FastDNA Spin Kit (MP, USA), and 500 ng DNA was used as the PCR template. 16S rRNA fragments of distinct regions (V4/V5) were amplified using the specific primers with Illumina adapter sequences (515F: GTGCCAGCMGCCGCGG and 907R: CCGTCAATTCMTTTRAGTTT). These PCR products were purified with a Gel Extraction Kit (Qiagen, Germany), and sequencing libraries were generated using TruSeq® DNA PCR-Free Sample Preparation Kit (Illumina, USA) following the manufacturer’s instructions. The library quality was assessed on the Qubit 3.0 (Thermo Fisher Scientific, USA) and Bioanalyzer 2100 system (Agilent, USA). Then the library was sequenced on an Illumina MiSeq platform and 2×250 bp paired-end reads were generated.

**Supplementary references**

1.	Marcia M, Ermler U, Peng GH & Michel H. A new structure-based classification of sulfide: quinone oxidoreductases. *Proteins* **78**, 1073-1083 (2010).

2.	Brito, J. A. et al. Structural and functional insights into sulfide: quinone oxidoreductase. *Biochemistry* **48**, 5613-5622 (2009).

3.	Kumar, S., Stecher, G., Li, M., Knyaz, C. & Tamura, K. MEGA X: Molecular evolutionary genetics analysis across computing platforms. *Mol Biol Evol* **35**, 1547-1549 (2018).

4.	Letunic, I. & Bork, P. Interactive tree of life (iTOL) v3: an online tool for the display and annotation of phylogenetic and other trees. *Nucleic Acids Res* **44**, W242-W245 (2016).

5.	Chen, Y. X. et al. SOAPnuke: a MapReduce acceleration-supported software for integrated quality control and preprocessing of high-throughput sequencing data. *Gigascience* **7** (2017).

6.	Delcher, A. L., Bratke, K. A., Powers, E. C. & Salzberg, S. L. Identifying bacterial genes and endosymbiont DNA with Glimmer. *Bioinformatics* **23**, 673-679 (2007).

**Supplementary Table S1. Chemical parameters of sites for SRB isolation and *in situ* cultivation**

| Chemical parameters | The site for isolation (2017) | The site for *in situ* cultivation (2020) |
| --- | --- | --- |
| Depth under the surface of ocean (m) | 1143 | 1146 |
| Longitude and latitude | 119º17'04.956''E, 22º06'58.384''N | 119º17'04.429''E, 22º07'01.523''N |
| pH | 7.68 | 7.59 |
| Temperature (°C) | 3.75 | 3.66 |
| CH_4_ (μM) | 2169 | 1723 |
| Sulfate (mM) | 28.3 | 28.1 |
| Dissolved oxygen (mg/L) | 3.07 | 3.25 |
| CO_2_ (PPM) | 812.34 | 815 |
| Salinity (‰) | 34.54 | 34.55 |

**Supplementary Table S2. Composition of trace elements solution**

| Components | Content |
| --- | --- |
| Na_2_EDTA·2H_2_O (pH 8.0) | 29 g |
| MnSO_4_·H_2_O | 3.26 g |
| CoCl_2_·6H_2_O | 1.8 g |
| ZnSO_4_·7H_2_O | 1.0 g |
| NiSO_4_·6H_2_O | 0.11 g |
| GuSO_4_·5H_2_O | 0.10 g |
| H_3_BO_3_ | 0.10 g |
| KAl(SO4)2·12H_2_O | 0.10 g |
| Na_2_MoO_4_·2H_2_O | 0.10 g |
| Na_2_WO_4_·2H_2_O | 0.10 g |
| Na_2_SeO_3_ | 0.05 g |
| Milli-Q water | 1 L |

**Supplementary Table S3. Composition of vitamins solution**

| Components | Content |
| --- | --- |
| Thiamine (VB1) | 50 mg |
| Riboflavin (VB2) | 50 mg |
| Nicotinic acid (VB3) | 50 mg |
| DL-Calcium pantothenate (VB5) | 50 mg |
| Pyridoxine hydrochloride (VB6) | 100 mg |
| Biotin (VB7) | 20 mg |
| Folic acid (VB9) | 20 mg |
| Cobalamin (VB12) | 1 mg |
| Para amino benzoic acid | 50 mg |
| Lipoic acid | 50 mg |
| Milli-Q water | 1 L |

**Supplementary Table S4. Strains and plasmids and primers used in this study**

|  | Description or sequence | Notes |
| --- | --- | --- |
| Strains |  |  |
| *Desulfovibrio* *marinus* CS1 | isolated from sediment in deep sea clod seep | This study |
| *Escherichia coli* DH5α | F^-^, φ80*lac*ZΔM15, Δ(*lac*ZYA-*arg*F)U169, *deo*R, *rec*A1, *end*A1, *hsd*R17(r_k_^-^, m_k_^+^), *pho*A *sup*E44, λ^-^, *thi*-1, *gyr*A96, *rel*A1 | Commercial |
| *E*. *coli* BL21(DE3) | F^-^, *omp*T, *hsd*S_B_(r_B_^-^, m_B_^-^), *gal*, *dcm*(DE3) | Commercial |
| Plasmids |  |  |
| pMD19-T simple | High copy number cloning vector, ^a^Amp^r^ | Commercial |
| pET28a (+) | Prokaryotic expression vector, ^a^Kan^r^ | Commercial |
| pET28a (+)/*sqr* | pET28a (+) containing the ORF^b^ of *sqr* gene, Kan^r^ | This study |
| pET28a (+)/*phsA* | pET28a (+) containing the ORF of *phsA* gene, Kan^r^ | This study |
| pET28a (+)/*sqr*-*phsA* | pET28a (+) containing both ORFs of *sqr* gene and *phsA* gene, Kan^r^ | This study |
| Primers |  |  |
| sqr-F | CGGGATCCATGCGCAAGATTCTGATCATCGG | *BamH*I |
| sqr-R | CCGAATTCAGGCTTTCCAGCGGCAAGTCCA | *EcoR*I |
| phsA-F | CGGAATTCATGAGCGATTTCAAAACTGTCTACA | *EcoR*I |
| phsA-R | CGGCTCGAGCGTCTTGCGCACTTCGATG | *Xho*I |
| sqr-RTF | TGCTGGTCTCCATCCCG |  |
| sqr-RTR | CCTGTGCCTTGAGCGTGT |  |
| phsA-RTF | CCGACAAGTGGATTCAGGAC |  |
| phsA-RTR | TGGATGCCAGATCACAGACG |  |
| 16S-RTF | CCCAATCACCAGCCCTACC |  |
| 16S-RTR | GGTCACGCCAATCCCAAA |  |

^a^ Amp^r^, ampicillin resistance; Kan^r^, kanamycin resistance.

^b^ ORF, open reading frame.

**Supplementary Table S5. Chemical parameters of the sampling sites for metagenomic sequencing and amplicon analysis**

| Chemical parameters for sample site | C1 | C4 | C2 | C3 | C5 |
| --- | --- | --- | --- | --- | --- |
| Depth under the surface of ocean (m) | 1139 | 1121 | 1137 | 1137 | 1137 |
| Depth under the surface of sediment (cm) | 0~20 | 20~40 | 40~60 | 120~140 | >280 |
| Longitude and latitude | E 119º17'09.106''  N 22º06'55.114'' | E 119º17'07.322''  N 22º06'58.598'' | E 119º17'06.436''  N 22º06'55.169'' | E 119º17'06.436''  N 22º06'55.169'' | E 119º17'06.436''  N 22º06'55.169'' |
| pH | 7.62 | NA | NA | NA | NA |
| Temperature (°C) | 3.64 | NA | NA | NA | NA |
| Salinity (‰) | 34.49 | NA | NA | NA | NA |
| Dissolved oxygen (mg/L) | 3.03 | NA | NA | NA | NA |
| CO_2_ (PPM) | 828 | NA | NA | NA | NA |
| CH_4_ (μM) | 2642 | NA | NA | NA | NA |
| Sulfate (mM) | 28 | NA | NA | NA | NA |
| S (PPM) | 3452.3 | 3644.6 | 5264.1 | 1860.1 | 3857.5 |

NA, not available.

**Supplementary Table S6. General features of *D*. *marinus* CS1 genome**

| *D*. *marinus* CS1 | Features |
| --- | --- |
| Accession number | CP039543.1 |
| Genome size (bp) | 4.76 M |
| G + C content (%) | 62.4% |
| Genes | 4169 |
| Proteins | 4089 |
| 5S rRNA genes | 2 |
| 16S rRNA genes | 2 |
| 23S rRNA genes | 2 |
| tRNA genes | 48 |
| Other RNAs | 4 |
| Pseudo Genes (total) | 21 |
| Chromosome number | 1 |

**Supplementary Table S7. Key proteins associated with sulfur metabolism**

**in *D. marinus* CS1**

| Locus_tag | Protein name | Nearest neighbor  and sequence |
| --- | --- | --- |
| E8L03_09635 | QmoC, quinone-interacting membrane -bound oxidoreductase complex subunit QmoC | *Desulfovibrio* *marinus*  WP_144233425.1 |
| E8L03_09640 | QmoB, hydrogenase iron-sulfur subunit | *Desulfovibrio* *marinus*  WP_171267232.1 |
| E8L03_09645 | QmoA, CoB-CoM heterodisulfide reductase iron-sulfur subunit A family protein | *Desulfovibrio* *marinus*  WP_144233423.1 |
| E8L03_09650 | AprA, adenylyl-sulfate reductase subunit alpha | *Desulfovibrio* *marinus*  WP_144233422.1 |
| E8L03_09655 | AprB, adenylyl-sulfate reductase subunit beta | *Desulfovibrio* *marinus*  WP_144233421.1 |
| E8L03_09660 | Sat, sulfate adenylyltransferase | *Desulfovibrio* *marinus*  WP_144233420.1 |
| E8L03_06435 | DsrC, TusE/DsrC/DsvC family sulfur relay protein | *Desulfovibrio* *marinus*  WP_144306202.1 |
| E8L03_15015 | DsrA, dissimilatory-type sulfite reductase subunit alpha | *Desulfovibrio* *marinus*  WP_144234054.1 |
| E8L03_15020 | DsrB, dissimilatory-type sulfite reductase subunit beta | *Desulfovibrio* *marinus*  WP_144234053.1 |
| E8L03_15025 | DsrD, dissimilatory sulfite reductase-asociated protein | *Desulfovibrio* *marinus*  WP_144234052.1 |
| E8L03_15030 | DsrN, cobyrinate a,c-diamide synthase | *Desulfovibrio* *marinus*  WP_171267775.1 |
| E8L03_20340 | DsrE, DsrE family protein | *Desulfovibrio* *marinus*  WP_144306673.1 |
| E8L03_20390 | DsrP, polysulfide reductase | *Desulfovibrio* *marinus*  WP_144306663.1 |
| E8L03_20395 | DsrO, 4Fe-4S dicluster domain -containing protein | *Desulfovibrio* *marinus*  WP_171268368.1 |
| E8L03_20400 | DsrJ, sulfate reduction electron transfer complex DsrMKJOP subunit | *Desulfovibrio* *marinus*  WP_144306661.1 |
| E8L03_20405 | DsrK, [DsrC]-trisulfide reductase | *Desulfovibrio* *alaskensis*  WP_011368154.1 |
| E8L03_20410 | DsrM, sulfate reduction electron transfer complex DsrMKJOP subunit | *Desulfovibrio* *marinus*  WP_144306659.1 |
| E8L03_20415 | DsrT, dissimilatory sulfite reductase system component | *Sulfobium* *mesophilum*  SPQ01222.1 |
| E8L03_05425 | SQR, sulfide: quinone oxidoreductase | *Syntrophobacter* sp. SbD2  WP_144305861.1 |
| E8L03_06385 | PhsA, thiosulfate reductase | *Desulfobulbus* sp. Tol-SR  KGO34598.1 |
| E8L03_06390 | DmsB, anaerobic dimethyl sulfoxide reductase subunit B | Desulfovibrionaceae bacterium KAF0234029.1 |
| E8L03_02130 | DsyB, MTHB methyltransferase; dimethylsulphoniopropionate biosynthesis enzyme | *Labrenzia* *aggregate*  AOR83342.1 |
| E8L03_10790 | SsuA, aliphatic sulfonate ABC transporter | *Desulfobulbus* *oralis*  AVD70971.1 |
| E8L03_08980 | CysC, adenylyl-sulfate kinase | *Desulfovibrio* *marinus*  WP_171267153.1 |
| E8L03_10175 | CysQ, 3'(2'),5'-bisphosphate nucleotidase | *Desulfovibrio* *marinus*  WP_171267284.1 |
| E8L03_16675 | CysH, phosphoadenosine phosphosulfate reductase family protein | *Desulfovibrio* *marinus*  WP_171267978.1 |
| E8L03_09290 | CysK, cysteine synthase | *Desulfovibrio* *marinus*  WP_171267194.1 |
| E8L03_07280 | MetB, cystathionine gamma-synthase family protein | *Desulfovibrio* *marinus*  WP_171266969.1 |
| E8L03_12345 | MetX, homoserine O-acetyltransferase | *Desulfovibrio* *marinus*  WP_171267515.1 |
| E8L03_19285 | MetE, 5-methyltetrahydropteroyltriglutamate -homocysteine S-methyltransferase | *Desulfovibrio* *marinus*  WP_171268249.1 |
| E8L03_08840 | MetH, methylenetetrahydrofolate reductase [NAD(P)H] | *Desulfovibrio* *marinus*  WP_144233579.1 |

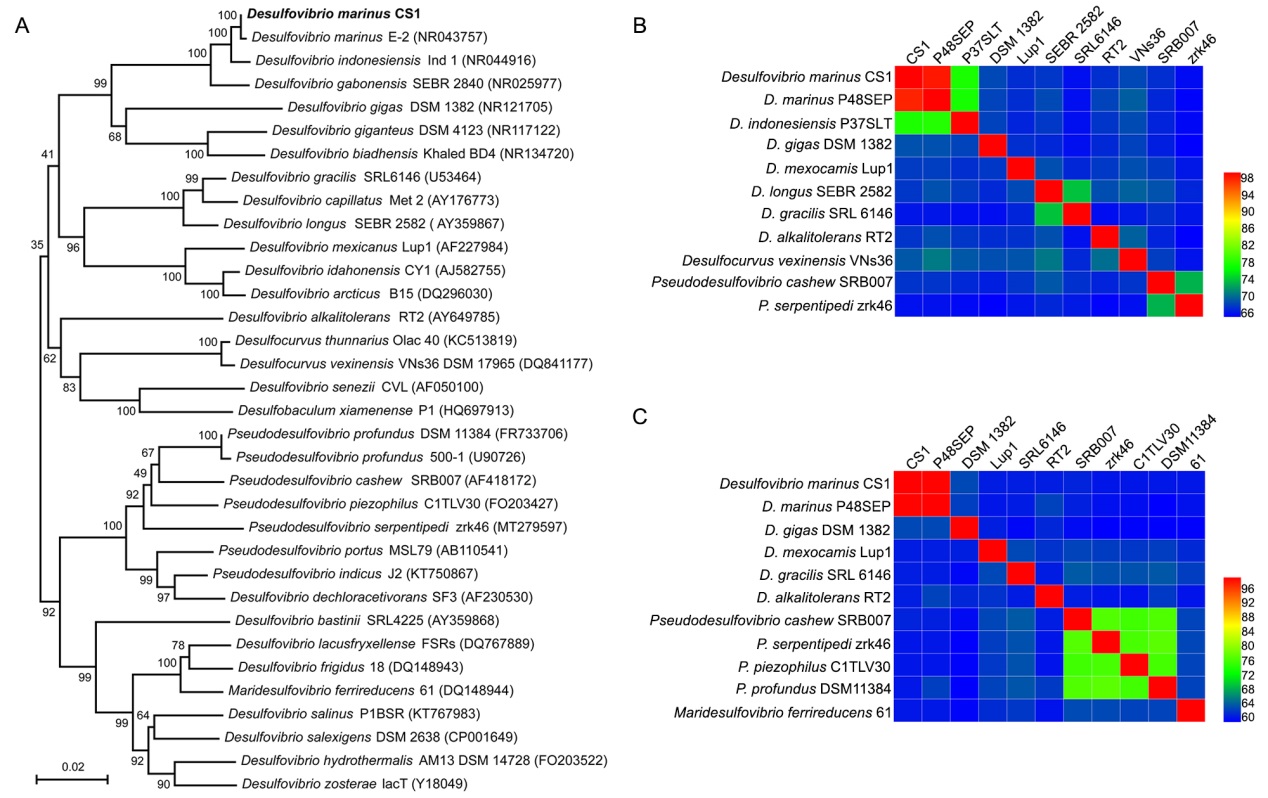
**Fig. S1** Phylogenetic analysis of *D*. *marinus* CS1. (A) Phylogenetic placement of strain CS1 within the representative species in the family Desulfovirbionaceae based on almost complete 16S rRNA gene sequences. The tree is inferred and reconstructed under the the neighbor-joining criterion and expressed as percentages of 1,000 replications. Bar, 0.02 substitutions per nucleotide position. (B-C) The ANI and AAI analyses of genomes between strain CS1 and other strains in the family Desulfovirbionaceae, respectively. The accepted threshold of ANI and AAI for same species is above 95%.

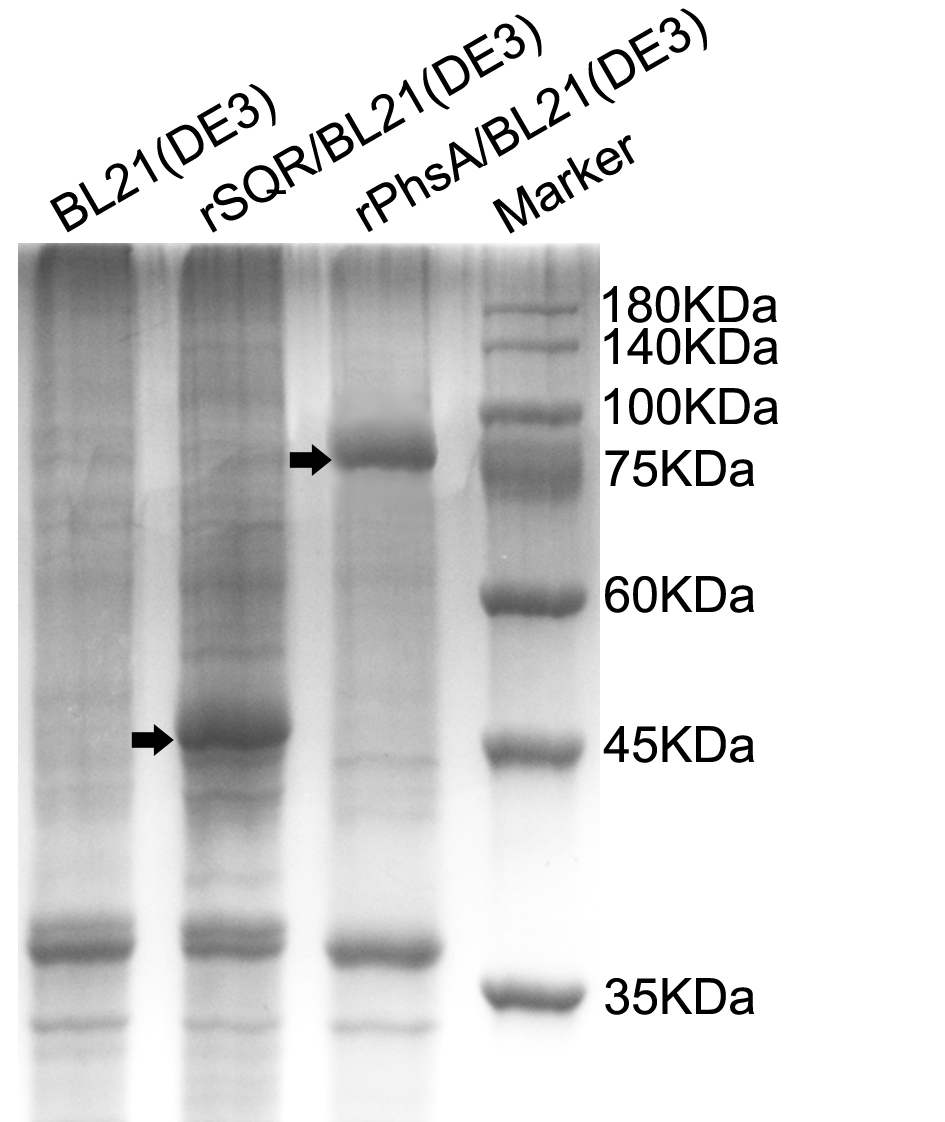

**Fig. S2** Detection of the overexpression of SQR and PhsA in *E*. *coli* BL21(DE3) via SDS-PAGE. Line 1, BL21(DE3) transformed with pET28a(+) as the negative control. Line 2, SQR of *D*. *marinus* CS1 was overexpressed in *E*. *coli* BL21(DE3). The arrow indicates the position of His-tagged SQR in the supernatant of *E*. *coli* *sqr/*pET28a (+)/BL21(DE3). Line 3, PhsA of *D*. *marinus* CS1 was overexpressed in *E*. *coli* BL21(DE3). The arrow indicates the position of His-tagged PhsA in the supernatant of *E*. *coli* *phsA/*pET28a (+)/BL21(DE3). Line 4, Standard molecular weight protein marker.

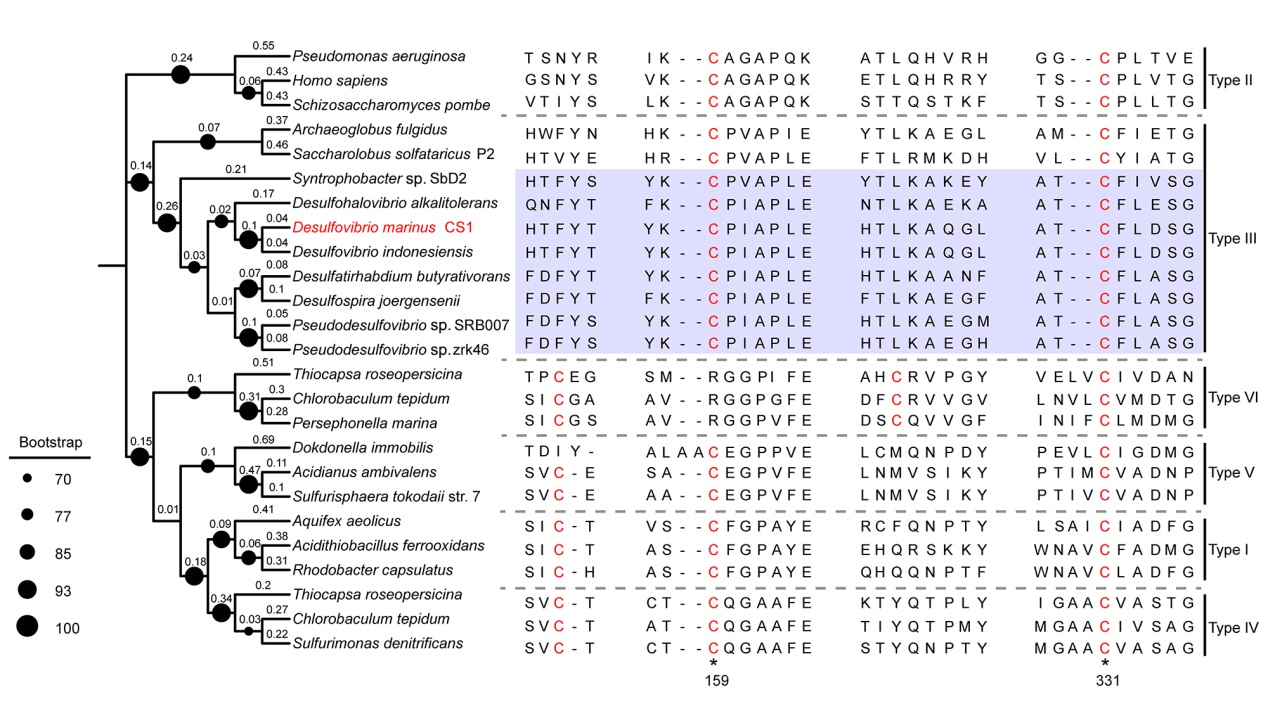

**Fig. S3** Phylogenetic analysis and sequence alignment of SQR homologs identified in *D. marinus* CS1 and other bacteria. The conserved sites of cysteine in different SQRs are highlighted with red color. The numbers at the bottom represent positions of two conserved cysteines in the SQR identified in *D. marinus* CS1. The tree is inferred and reconstructed under the maximum likelihood criterion, and bootstrap values (%) > 70 are indicated at the base of each node (expressed as black nots with different sizes). Percentages represent the tree scale of each branch.

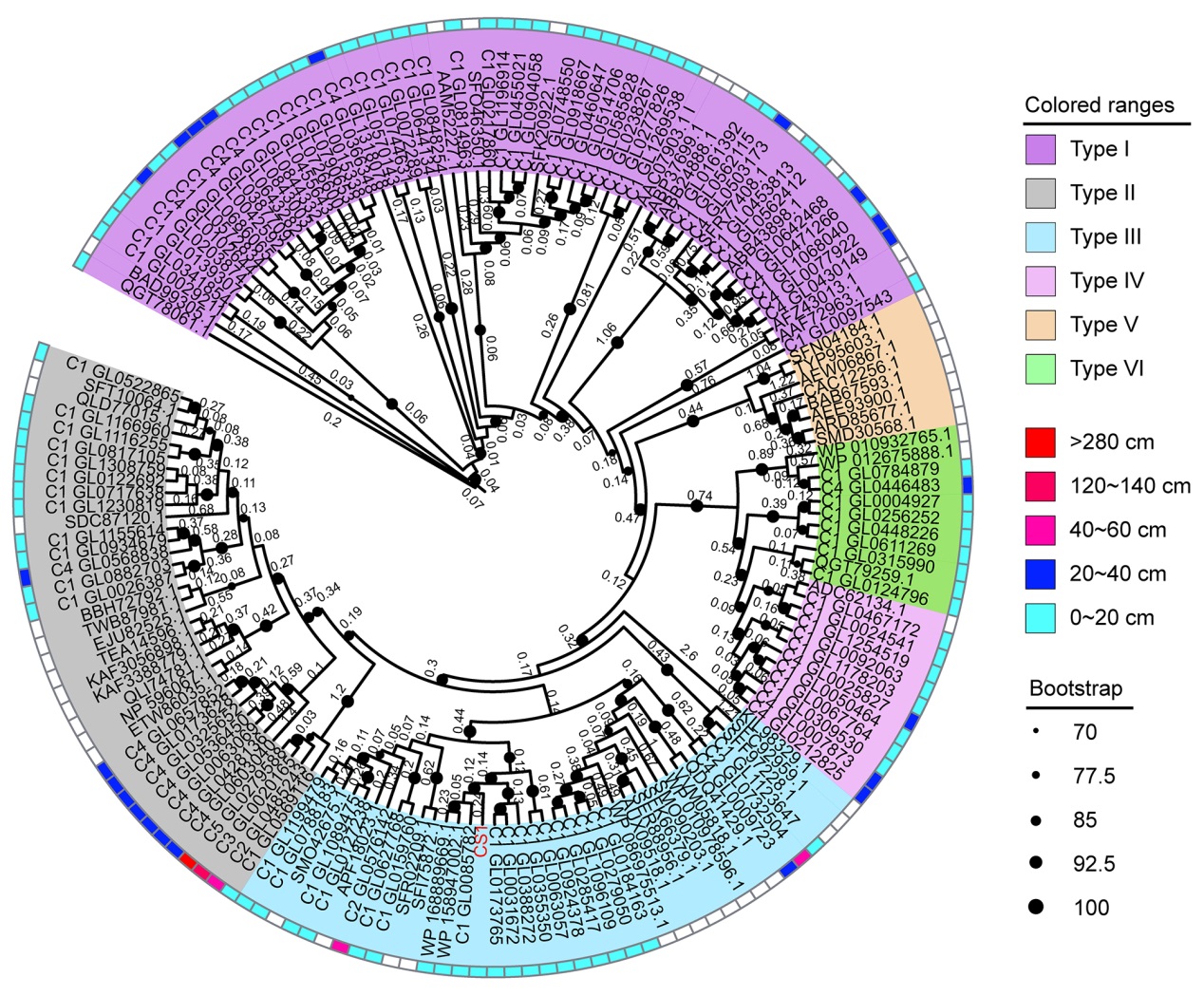

**Fig. S4** Maximum-likelihood phylogeny of sulfide:quinone oxidoreductases (SQRs) identified in metagenomic data derived from deep-sea cold seep sediments (1,000 bootstrap replicates). Bootstrap values (%) > 70 are indicated at the base of each node (expressed as black nots with different sizes). Percentages represent the tree scale of each branch. The detailed information of proteins used for phylogenetic analyses is listed in the Supplementary Dataset 1.
